# Control of brown adipose tissue adaptation to nutrient stress by the activin receptor ALK7

**DOI:** 10.1101/861609

**Authors:** Patricia Marmol-Carrasco, Carlos F. Ibáñez

## Abstract

Adaptation to nutrient availability is crucial for survival. Upon nutritional stress, such as during prolonged fasting or cold exposure, organisms need to balance the feeding of tissues and the maintenance of body temperature. The mechanisms that regulate the adaptation of brown adipose tissue (BAT), a key organ for non-shivering thermogenesis, to variations in nutritional state are not known. Here we report that specific deletion of the activin receptor ALK7 in BAT resulted in fasting-induced hypothermia due to exaggerated catabolic activity in brown adipocytes. After overnight fasting, BAT lacking ALK7 showed increased expression of genes responsive to nutrient stress, including the upstream regulator KLF15, aminoacid catabolizing enzymes, notably proline dehydrogenase (POX), and adipose triglyceride lipase (ATGL), as well as markedly reduced lipid droplet size. In agreement with this, ligand stimulation of ALK7 suppressed POX and KLF15 expression in both mouse and human brown adipocytes. Treatment of mutant mice with the glucocorticoid receptor (GR) antagonist RU486 restored KLF15 and POX expression levels in mutant BAT, suggesting that loss of BAT ALK7 results in excessive activation of glucocorticoid signaling upon fasting. These results reveal a novel signaling pathway downstream of ALK7 which regulates the adaptation of BAT to nutrient availability by limiting nutrient stress-induced overactivation of catabolic responses in brown adipocytes.

## Introduction

The adipose depots in mammals consist mainly of white (WAT) and brown (BAT) adipose tissues (Frontini and Cinti, 2010; Rosen and Spiegelman, 2014). WAT stores energy in the form of triglycerides, which can be mobilized in time of higher energy expenditure or nutrient scarcity. Fat accumulation in WAT is an anabolic process, mainly regulated by insulin, while fat breakdown by lipolysis can be considered as catabolic and is predominantly controlled by catecholamines. The mitochondria of BAT contain Uncoupled Protein 1 (UCP1) and defends body temperature against low environmental temperature producing heat through non-shivering thermogenesis (Cannon and Nedergaard, 2004). Under certain conditions, subcutaneous WAT depots can also develop UCP1-expressing cells, referred to as beige or brite adipocytes (Wu et al., 2013; Nedergaard and Cannon, 2014). Cold sensing in mammals results in the transduction of a signal from the central nervous system to the sympathetic nerve endings in BAT. Adrenergic stimulation in BAT triggers the release of long-chain fatty acids from cytoplasmic lipid droplets (Cannon and Nedergaard, 2004), which act as both the main energy substrate for thermogenesis as well as activators of H^+^ transport activity in UCP1 (Fedorenko et al., 2012). Dissipation of the mitochondrial H^+^ gradient generated during substrate oxidation results in heat production at the expense of ATP synthesis (Nicholls and Locke, 1984). Other origins of fatty acids for thermogenesis include WAT lipolysis as well as dietary fat. During non-shivering thermogenesis, cytosolic glucose oxidation is also greatly increased to counterbalance the decrease in ATP synthesis that results from uncoupling (Hao et al., 2015). While mainly driven by the central nervous system, non-shivering thermogenesis is also known to be regulated by secreted factors (Villarroya and Vidal-Puig, 2013).

Due to the high amounts of energy consumed by BAT, non-shivering thermogenesis is highly sensitive to nutrient availability and functions optimally during well-fed conditions. In periods of nutrient scarcity, mammals are able to reduce thermogenesis as an adaptive, energy-saving mechanism by entering torpor, a state characterized by decreased activity and body temperatures lower than 32°C (Geiser et al., 2014). Mice can enter torpor when confronted with severe nutrient stress, such as during prolonged fasting, and can be exacerbated by cold exposure (Oelkrug et al., 2011). During a situation of nutrient stress combined with low ambient temperature, animals must reconcile the requirements of high energy demanding organs, such as the brain, with the need to maintain body temperature. However, our understanding of the mechanisms that control this balance is limited. Mechanisms underlying responses to nutritional stress are better understood in liver and muscle, and include the induction by glucocorticoids of Kruppel Like Factor 15 (KLF15), a key regulator of nutritional adaptations during fasting, such as liver gluconeogenesis and amino acid catabolism in muscle (Yamamoto et al., 2004; Gray et al., 2007; Shimizu et al., 2011). Whether BAT employs similar or different mechanisms to adapt its thermogenic activity to fluctuations in nutrient availability is currently unknown.

ALK7, encoded by the *Acvr1c* gene, is a type I receptor of the TGF-β receptor superfamily that mediates the activities of a diverse group of ligands, including activin B, growth and differentiation factor 3 (GDF-3) and Nodal (Rydén et al., 1996; Reissmann et al., 2001; Andersson et al., 2008). In rodents as well as humans, ALK7 is expression is enriched in tissues that are important for the regulation of energy homeostasis, including adipose tissue (Andersson et al., 2008), pancreatic islets (Bertolino et al., 2008), endocrine gut cells (Jörnvall et al., 2004) and the arcuate nucleus of the hypothalamus (Sandoval-Guzmán et al., 2012). In white adipose tissue (WAT), previous studies have shown that ALK7 signaling facilitates fat accumulation by repressing the expression of adrenergic receptors, thereby reducing catecholamine sensitivity and lipolysis (Guo et al., 2014). Accordingly, mutant mice lacking ALK7 globally, or only in adipocytes, are resistant to diet-induced obesity (Andersson et al., 2008; Guo et al., 2014). Recent studies have identified polymorphic variants in the human *Acvr1c* gene which affect body fat distribution and protect from type II diabetes (Emdin et al., 2019; Justice et al., 2019), indicating that ALK7 has very similar functions in humans as in rodents. The function of ALK7 in BAT has not been investigated, and the present study represents the first attempt to understand its role in BAT physiology.

## Results

### Fasting induces abnormally increased fat catabolism in BAT of *Ucp1*^CRE^:*Alk7*^fx/fx^ mutant mice lacking ALK7 in brown adipocytes

Expression of *Alk7* mRNA was detected in interscapular BAT (iBAT) of young adult male mice at levels comparable to those found in inguinal WAT (iWAT), although lower than *Alk7* mRNA expression in epididymal WAT (eWAT) (Figure 1A). No *Alk7* mRNA expression could be detected in liver. The level of *Alk7* mRNA was low in cells isolated from BAT stromal vascular fraction (SVF), containing precursors of brown adipocytes, but was markedly upregulated after *in vitro* differentiation into brown adipocytes, reaching levels comparable to those found in mature adipocytes freshly isolated from BAT (Figure 1B). In order to investigate cell-autonomous functions of ALK7 in BAT, we generated mice lacking this receptor specifically in brown adipocytes by crossing *Alk7*^fx/fx^ mice (Guo et al., 2014) with *Ucp1*^CRE^ mice (Kong et al., 2014), expressing CRE recombinase under regulatory sequences of the gene encoding Uncoupling Protein 1 (*Ucp1*). In the resulting mutant mice (*Ucp1*^CRE^:*Alk7*^fx/fx^), *Alk7* mRNA expression in BAT was almost completely abolished (Figure 1C), confirming that ALK7 is exclusively expressed by brown adipocytes in BAT. At 2 months of age, *Ucp1*^CRE^:*Alk7*^fx/fx^ mutant mice showed body weight and fat composition indistinguishable from control mice under Chow diet (Figure 1D). Energy consumption was not affected by the lack of ALK7 in BAT, as demonstrated by normal oxygen consumption (VO_2_) under both fed and fasted conditions (Figure S1A). Similarly, respiratory exchange ratio (RER), which reports the relative usage of carbohydrates and fat, and food intake were also comparable between *Ucp1*^CRE^:*Alk7*^fx/fx^ and control mice (Figures S1B and C). Both iBAT and eWAT mass relative to body weight were also normal in 2 month old mutants (Figure 1E), as well as expression of a battery of BAT differentiation and maturation markers and mitochondrial copy number (Figures S2A and B), indicating that ALK7 is not required by brown adipocytes for normal BAT development. Interestingly, overnight (14h) fasting also induced a significant reduction in iBAT weight in 2 month old *Ucp1*^CRE^:*Alk7*^fx/fx^ mice, despite normal eWAT mass (Figure 1F).

**Figure 1.**
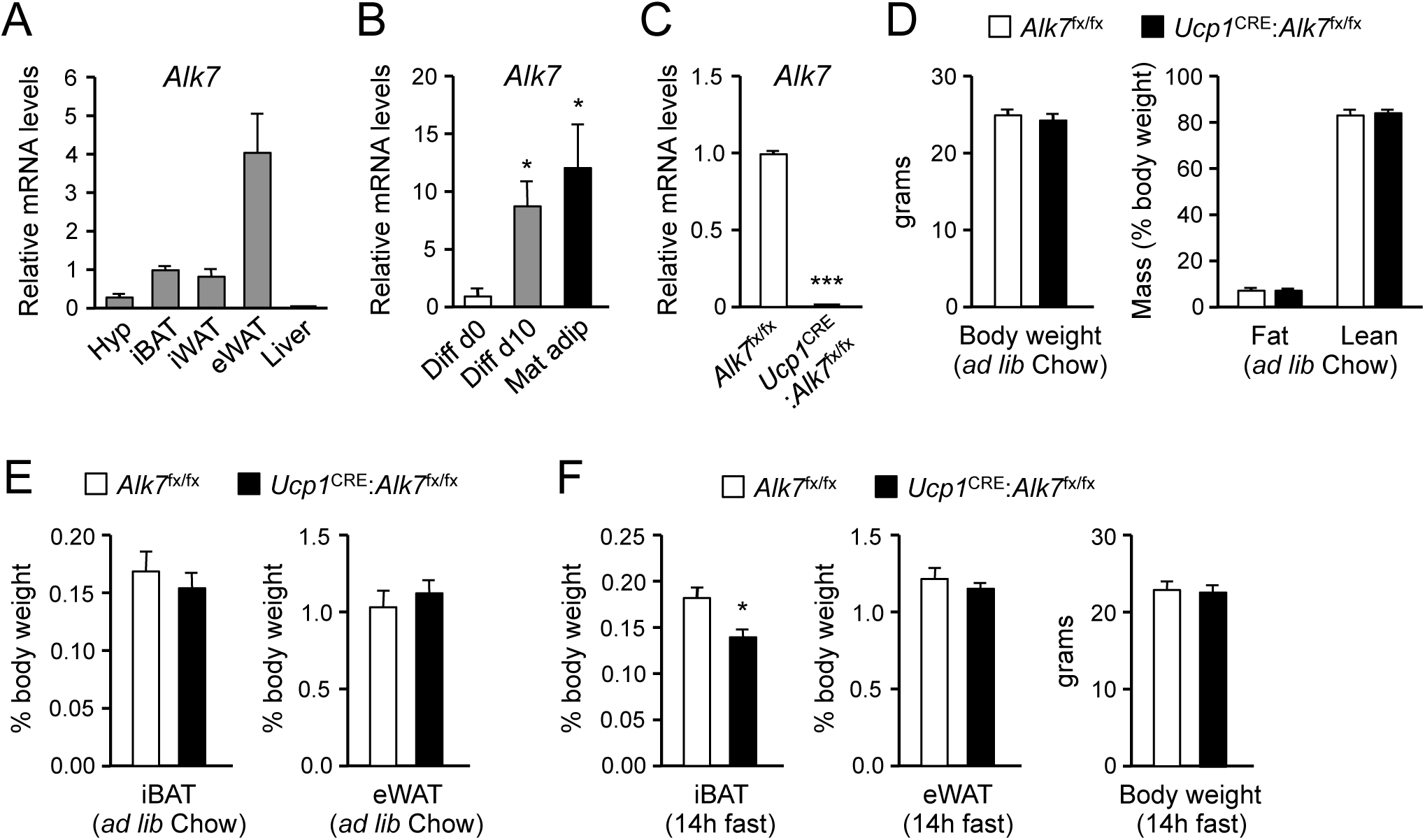
Reduced iBAT mass in *Ucp1*^CRE^:*Alk7*^fx/fx^ conditional mutant mice after fasting. (A) Q-PCR determination of *Alk7* mRNA expression in hypothalamus (Hyp), interscapular BAT (iBAT), inguinal WAT (iWAT), epididymal WAT (eWAT) and liver of wild type mice. The values were normalized to mRNA levels in iBAT and are presented as average ± SEM. N=4 or 6 (iWAT) mice per group. (B) Q-PCR determination of *Alk7* mRNA expression in iBAT stromal vascular fraction (Diff d0), adipocytes differentiated *in vitro* (Diff d10), and freshly isolated mature adipocytes (Mat adip). The values were normalized to mRNA levels in the Diff d0 sample, and are presented as average ± SEM. N=4 independent experiments. *, P<0.05; two-tailed Mann Whitney test. (C) Q-PCR determination of *Alk7* mRNA expression in iBAT from conditional mutant (*Ucp1*^CRE^:*Alk7*^fx/fx^) and control (*Alk7*^fx/fx^) mice. The values were normalized to mRNA levels in control mice and are presented as average ± SEM. N=4 mice per group. ***, P<0.001; two-tailed Mann Whitney test. (D) Body weight (left) at 2 months (*ad libitum* Chow diet) and fat and lean mass (expressed as percentage of body weight) assessed by magnetic resonance imaging (right). Values are presented as average ± SEM. N=4 mice per group. (E) Relative iBAT and eWAT mass expressed as percentage of body weight at 2 months (*ad libitum* Chow diet) in conditional mutant and control mice. Values are presented as average ± SEM. N=5 mice per group. (F) Relative iBAT mass, eWAT mass, and body weight at 2 months (*ad libitum* Chow diet) following 14h fasting in conditional mutant and control mice. Values are presented as average ± SEM. N=5 mice per group. *, P<0.05; two-tailed unpaired Student’s t-test.

Histological analysis of iBAT revealed a significant decrease in lipid droplet size in 2 month old *Ucp1*^CRE^:*Alk7*^fx/fx^ mice compared to age-matched controls (Figure 2A). A similar reduction was observed in the iBAT of global *Alk7*^-/-^ knock-out mice (Figure 2B). A proteomics analysis of iBAT lipid droplets from *Alk7*^-/-^ knock-out mice revealed increased levels of adipose triglyceride lipase (ATGL), the rate-limiting enzyme of lipolysis, despite normal levels of other major lipid droplet proteins, including hormone-sensitive lipase (HSL) (Figure 2C). Total iBAT lysates from *Ucp1*^CRE^:*Alk7*^fx/fx^ conditional mutant mice showed increased ATGL levels compared to controls (Figure 2D), although the difference did not reach statistical significance (P=0.077). However, ATGL protein levels were robustly increased in the mutant iBAT 14h after fasting (Figure 2E), which was in line with the reduced iBAT mass observed in fasted mutant animals (Figure 1H). ATGL levels were not changed by fasting in BAT of wild type mice (Figure S3A). No differences could be detected in the levels of phosphorylated HSL (P-HSL^S563^, Figures 2D and E). Enhanced ATGL protein levels in the mutants was accompanied by a strong trend (P=0.078) towards increased *Atgl* mRNA levels after fasting (Figure 2F). iBAT from fasted conditional mutant mice also showed a significant decrease in the mRNA levels of G0/G1 switch gene 2 (*G0S2*), which encodes an inhibitor of ATGL activity (Yang et al., 2010) (Figure 2G). These changes in ATGL and G0S2 expression prompted us to examine basal lipolysis in iBAT explants from *Ucp1*^CRE^:*Alk7*^fx/fx^ and control mice as measured by basal glycerol release. Significantly increased levels of glycerol were detected in explant supernatants derived from fasted mutant mice compared to controls (Figure 2H), indicating abnormally enhanced basal lipolysis in iBAT of *Ucp1*^CRE^:*Alk7*^fx/fx^ mice after fasting.

**Figure 2.**
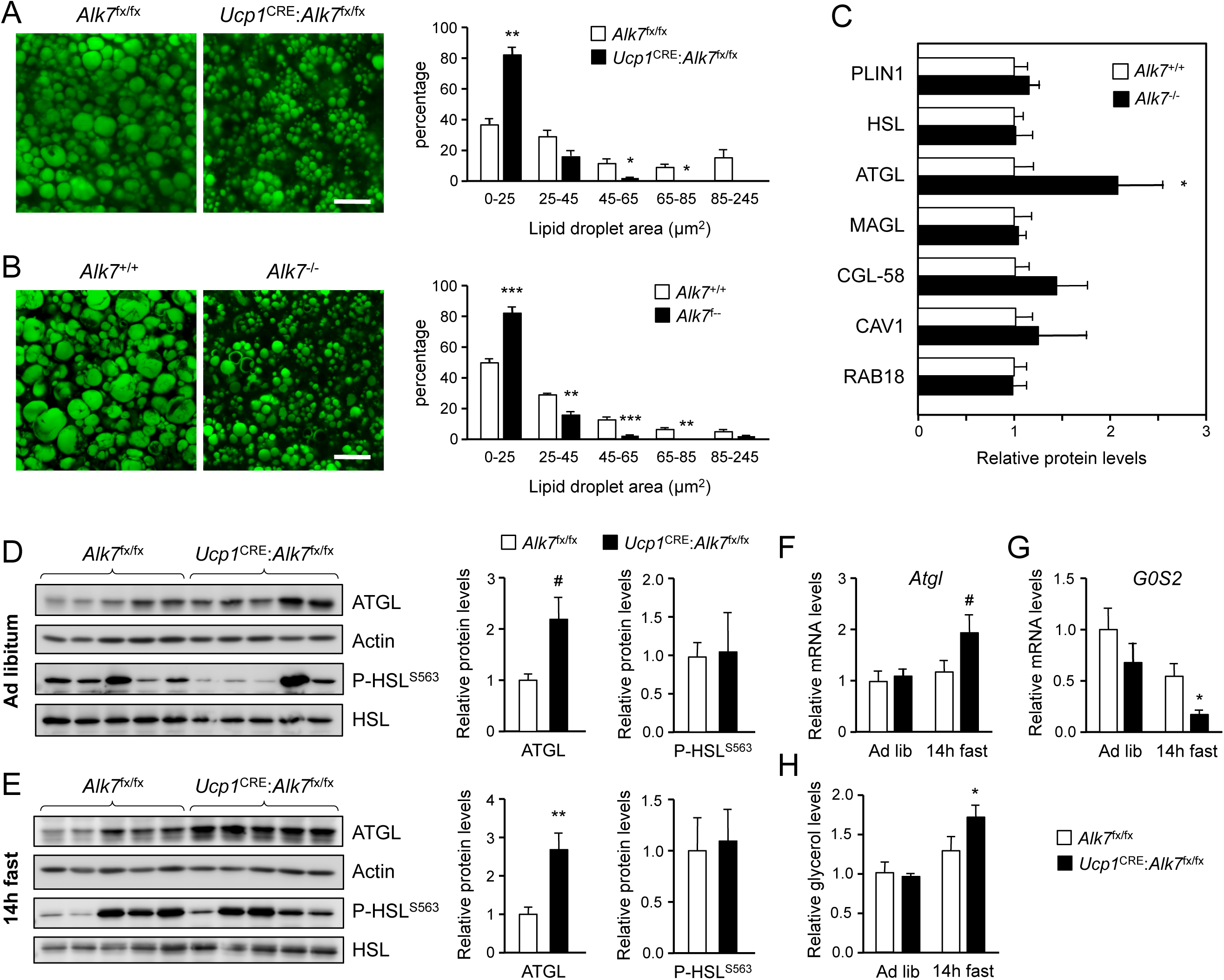
Fasting induces abnormally increased fat catabolism in BAT of *Ucp1*^CRE^:*Alk7*^fx/fx^ conditional mutant mice. (A, B) Representative BODIPY 493/503 staining of iBAT sections of conditional mutant (*Ucp1*^CRE^:*Alk7*^fx/fx^) and control (*Alk7*^fx/fx^) mice (A) and global *Alk7*^-*/*-^ knock-out mice and wild type controls (B). Scale bar, 20 µm. Histograms to the right show quantitative analysis of lipid droplet size. Values are presented as average ± SEM. N=4 mice per group. *, P<0.05; **, P<0.01; ***, P<0.001; vs. control, respectively; unpaired Student’s t-test. (C) Proteomic analysis of BAT lipid droplet fractions in global *Alk7*^-*/*-^ knock-out mice and wild type controls. PLIN1, Perilipin 1; HSL, Hormone-sensitive lipase; ATGL, Adipose triglyceride lipase; MAGL, Monoacylglycerol lipase; CGL-58, Comparative Gene Identification-58; CAV1, caveolin-1. Values are presented as average ± SEM. N=4 mice per genotype. *, P<0.05, unpaired Student’s t-test. (D, E) Western blot analysis of ATGL and phosphorylated HSL (P-HSL^S563^) in iBAT of 2 month old conditional mutant and control mice fed Chow *ad libitum* (D) or after 14h fasting (E). Histograms to the right show quantitative analyses of protein levels normalized to those in control *Alk7*^fx/fx^ mice. N=5 mice per genotype. #, P=0.077; *, P<0.05; two-tailed unpaired Student’s t-test. (F, G) Q-PCR determination of *Atgl* (F) and *G0S2* (G) mRNA expression in iBAT of 2 month old conditional mutant and control mice fed Chow *ad libitum* or after 14h fasting. The values were normalized to mRNA levels in control mice fed *ad libitum* and are presented as average ± SEM. N=4 mice per group. #, P=0.078 vs. control; *, P<0.05 vs. control; two-tailed unpaired Student’s t-test. (H) Basal lipolysis measured as glycerol release *ex vivo* from iBAT explants from conditional mutant and control mice fed *ad libitum* or after 6h fasting. Values were normalized to *ad libitum* levels in control mice and are presented as average ± SEM. N=6 (ad lib) and 5 (fast) mice per group. *, P<0.05 vs. control; two-tailed unpaired Student’s t-test.

### Abnormally enhanced amino acid catabolism upon nutrient stress in iBAT lacking ALK7

The enhanced fat catabolism observed in iBAT lacking ALK7, particularly under fasting conditions, prompted us to investigate pathways involved in the regulation of metabolic balance. Insulin is a well known negative regulator of catabolic activity in adipose tissue during a postprandial state. Basal levels of AKT, a key downstream effector of insulin signaling, phosphorylated on Ser^473^ (P-AKT_S473_), which correlates with its activation state, were unchanged in iBAT of conditional mutant mice relative to controls, both under fed or fasted conditions (Figure S3B). P-AKT_S473_ levels were not changed by fasting in BAT of wild type mice (Figure S3C). In addition, P-AKT_S473_ levels were increased to the same extent in iBAT of both mutant and control mice in response to acute insulin administration (Figure S3D), indicating normal insulin sensitivity in iBAT lacking ALK7. A microarray analysis of iBAT from *Alk7*^-/-^ global knock-out mice revealed several changes in gene expression in mutant iBAT compared to wild type controls (Figures S4A and B), including a marked upregulation in the level of *Prodh* mRNA, encoding proline dehydrogenase (POX), a mitochondrial enzyme that catalyzes the first step in the degradation of proline, and a critical component of metabolic responses to nutrient stress in cancer cells (Pandhare et al., 2009; Phang, 2019). In line with this, *Prodh* mRNA levels were increased after 14h fasting in iBAT of both *Ucp1*^CRE^:*Alk7*^fx/fx^ conditional mutant and control mice, but significantly more so in the mutants (Figure 3A). Similarly, mRNAs encoding enzymes involved in the degradation of alanine and branched amino acids, namely ALT1 and BCAT2, were also specifically upregulated upon fasting in the mutant iBAT (Figure 3B). Realizing that these genes are all targets of KLF15, a key regulator of responses to nutritional stress in liver and skeletal muscle (Gray et al., 2007; Haldar et al., 2012), we examined the levels of *Klf15* mRNA in mutant and control iBAT. This revealed a significant induction of *Klf15* mRNA after fasting in iBAT from *Ucp1*^CRE^:*Alk7*^fx/fx^ mutant mice, but not from control mice (Figure 3C). As expected, fasting induced *Klf15* mRNA in the liver regardless of genotype (Figure 3C). *Klf15* is one major target of glucocorticoid signaling, mediating several catabolic responses to glucocorticoids in liver and skeletal muscle (Shimizu et al., 2011; Sasse et al., 2013), but its role and regulation in BAT had not been investigated. We first verified that overnight fasting increased corticosterone levels in serum of both *Ucp1*^CRE^:*Alk7*^fx/fx^ mutants and control mice to a similar extent (Figure 3D). We then asked whether *Klf15* mRNA expression in iBAT was similarly sensitive to glucocorticoid signaling, as it has been demonstrated in other tissues. To this end, we administered the glucocorticoid receptor antagonist RU486 to conditional mutant and control mice 4h prior to the end of a 14h fasting period, after which iBAT was collected for mRNA analysis. *Klf15* mRNA levels were reduced by RU486 treatment in both strains of mice (Figure 3E), indicating that *Klf15* expression is also under glucocorticoid regulation in BAT. Importantly, RU486 treatment also restored *Prodh* mRNA levels in iBAT of conditional mutant mice back to the level found in control mice (Figure 3F). Expression of *Alt1* and *Bcat2* mRNAs were not affected by RU486 treatment (Figure 3G).

**Figure 3.**
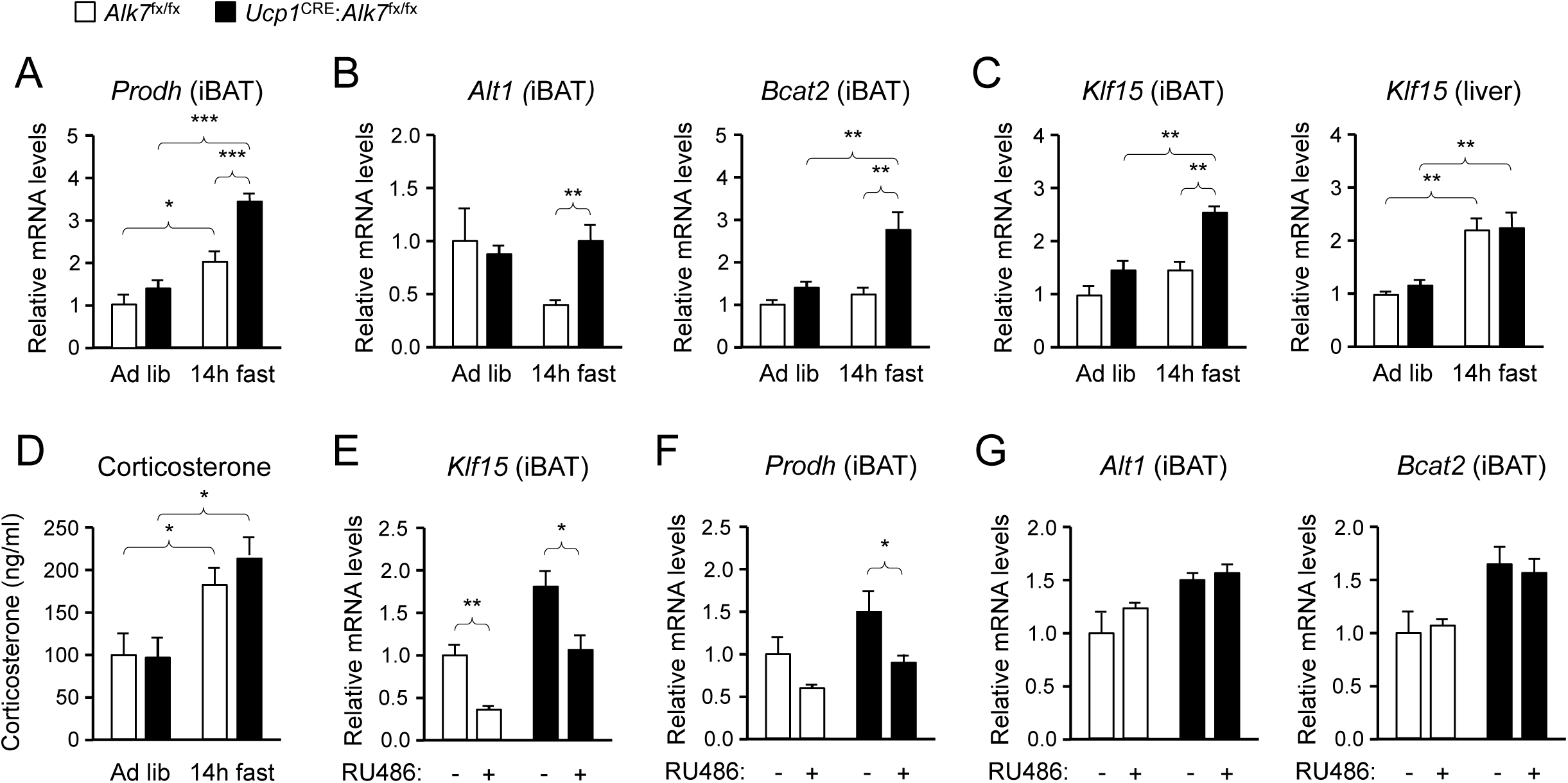
Abnormally enhanced amino acid catabolism upon nutrient stress in iBAT lacking ALK7. (A, B) Q-PCR determination of *Prodh* (A) and *Alt1* and *Bcat2* (B) mRNA expression in iBAT of 2 month old conditional mutant and control mice fed Chow *ad libitum* or after 14h fasting. The values were normalized to mRNA levels in control mice fed *ad libitum* and are presented as average ± SEM. N=9 (A) or 5 (B) mice per group. *, P<0.05; ***, P<0.001; vs. control, respectively; unpaired Student’s t-test. (C) Q-PCR determination of *Klf15* mRNA expression in iBAT (left) and liver (right) of 2 month old conditional mutant and control mice fed Chow *ad libitum* or after 14h fasting. The values were normalized to mRNA levels in control mice fed *ad libitum* and are presented as average ± SEM. N=5 mice per group. **, P<0.01; vs. control, respectively; two-way ANOVA with Bonferroni post-test. (D) Serum corticosterone levels (ng/ml) in 2 month old conditional mutant and control mice fed Chow *ad libitum* or after 14h fasting. Values are presented as average ± SEM. N=9 mice per group. *, P<0.05 vs. control; unpaired Student’s t-test. (E-G) Q-PCR determination of *Klf15* (E), *Prodh* (F) and *Alt1* and *Bcat2* (G) mRNA expression in iBAT of 10h-fasted conditional mutant and control mice 4h after injection with RU486 or vehicle, as indicated. The values were normalized to mRNA levels in control mice injected with vehicle, and are presented as average ± SEM. N=5 mice per group. *, P<0.05; **, P<0.01; vs. control, respectively; two-way ANOVA with Bonferroni post-test.

### Activin B suppresses expression of mRNAs encoding KLF15 and amino acid degrading enzymes in isolated mouse and human brown adipocytes

The results described above indicated that ALK7 signaling may negatively regulate the expression of genes involved in lipid and amino acid catabolism in brown adipocytes. In order to test this more directly, we established cultures of brown adipocytes derived by differentiation *in vitro* of iBAT SVF extracted from 2 month old wild type mice. After 10 days of differentiation, the expression of mRNAs for *Klf15*, *Prodh*, *Alt1* and *Bcat2* were strongly upregulated in these cultures compared to the levels in iBAT SVF cells (Figure 4A). Treatment with the ALK7 ligand activin B significantly reduced the expression of the four mRNAs in differentiated brown adipocytes (Figure 4B). This response was effectively suppressed by SB431542, an inhibitor of type I receptors for TGF-βs and activins, including ALK7 (Inman et al., 2002). Interestingly, SB431542 could on its own moderately increase the mRNA expression of the four genes, although the effect did not reach statistical significance (Figure 4B), perhaps reflecting the activities of endogenously produced ligands. Activin B had no effect on brown adipocytes lacking ALK7 (Figure 4C), indicating that its effects were mediated by the ALK7 receptor. In line with the reduced *G0S2* mRNA levels observed in iBAT from fasted *Ucp1*^CRE^:*Alk7*^fx/fx^ mutants (Figure 2F), treatment with activin B increased, while SB431542 decreased, expression of this gene in cultured brown adipocytes (Figure 4D). We detected a trend towards reduction of *Atgl* mRNA expression with activin B, and increased expression with SB431542, in agreement with the results *in vivo*, but these trends did not reach statistical significance. Regulation of *Atgl* mRNA expression by ALK7 may require additional, fasting-induced signals. Activin B had not significant effects on the mRNA levels of *Ucp1* or *Prdm16* (Figure S5). Importantly, similar observations could be made in cultured human brown adipocytes, in which activin B treatment also resulted in reduced levels of *Klf15* and *Prodh* mRNAs (Figure 4E). Together these results suggest that ALK7 signaling can directly suppress the expression of a series of mRNAs encoding diverse regulators of fat and amino acid catabolism.

**Figure 4.**
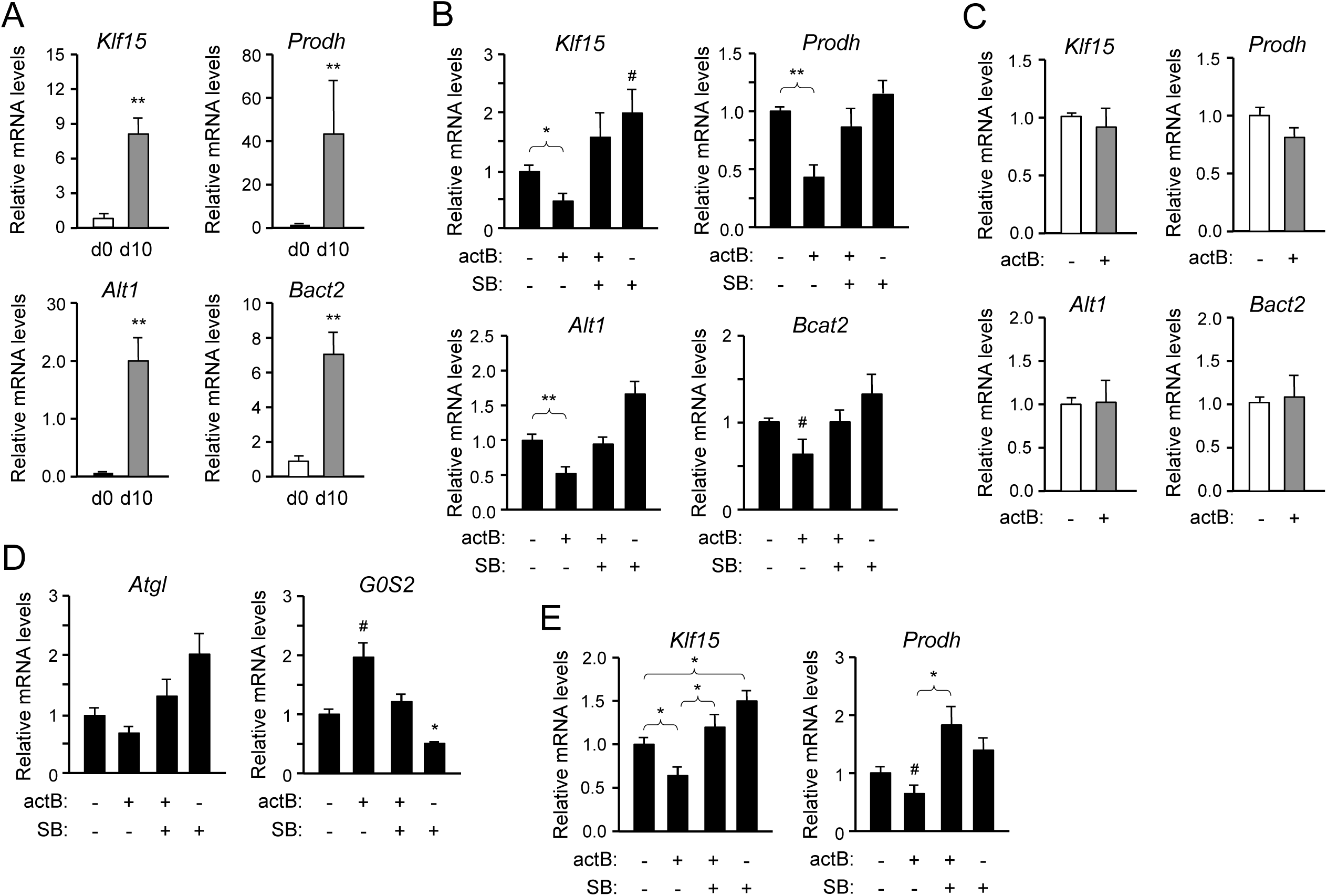
Activin B suppresses expression of mRNAs encoding KLF15 and amino acid degrading enzymes in isolated mouse and human brown adipocytes. (A) Q-PCR determination of *Klf15*, *Prodh*, *Alt1* and *Bcat2* mRNA expression in primary cultures of differentiated (d10) compared to non-differentiated (d0) brown adipocytes isolated from iBAT of wild type mice. The values were normalized to mRNA levels at d0 and are presented as average ± SEM. N=5 independent experiments each performed in duplicate. **, P<0.01 vs. d0; unpaired Student’s t-test. (B) Q-PCR determination of *Klf15*, *Prodh*, *Alt1* and *Bcat2* mRNA expression in primary cultures of differentiated brown adipocytes isolated from iBAT of wild type mice treated with activin B (actB) or SB-431542 (SB) for 24h as indicated. The values were normalized to levels in untreated cultures and are presented as average ± SEM. N=5 independent experiments each performed in duplicate. #, P=0.088; *, P<0.05; **, P<0.01 vs. untreated; two-tailed paired Student’s t-test. (C) Q-PCR determination of *Klf15*, *Prodh*, *Alt1* and *Bcat2* mRNA expression in primary cultures of differentiated brown adipocytes isolated from iBAT of *Alk7*^-*/*-^ knock-out mice treated with activin B (actB) for 24h as indicated. The values were normalized to levels in untreated cultures and are presented as average ± SEM. N=3 independent experiments each performed in duplicate. (D) Q-PCR determination of *Atgl* and *G0S2* mRNA expression in primary cultures of differentiated brown adipocytes isolated from iBAT of wild type mice treated with activin B (actB) or SB-431542 (SB) for 24h as indicated. The values were normalized to levels in untreated cultures and are presented as average ± SEM. N=5 independent experiments each performed in duplicate. #, P=0.05; *, P<0.05 vs. untreated; two-tailed paired Student’s t-test. (E) Q-PCR determination of *Klf15* and *Prodh* mRNA expression in primary cultures of differentiated human brown adipocytes isolated treated with activin B (actB) or SB-431542 (SB) for 24h as indicated. The values were normalized to levels in untreated cultures and are presented as average ± SEM. N=5 independent experiments each performed in duplicate. *, P<0.05; vs. untreated; two-tailed unpaired Student’s t-test.

### Increased proline-dependent ATP generation in mitochondria from iBAT lacking ALK7

The increased levels of *Prodh* mRNA, encoding the mitochondrial enzyme POX, in mutant iBAT prompted us to examine the levels of a battery of mitochondrial proteins, including POX, UCP1 and components of respiratory complexes I to V, in iBAT from *Ucp1*^CRE^:*Alk7*^fx/fx^ mutant and control mice, fed ad libitum as well as after 14h fasting. iBAT from mutant mice showed a moderate increase in POX protein levels compared to control mice (Figure 5A). However, fasting induced a significantly greater increase in POX protein levels in the mutant iBAT (Figure 5B). In addition, SDHA, a subunit of succinate dehydrogenase, was also differentially enhanced by fasting in the mutant iBAT (Figure 5B). Interestingly, POX and succinate dehydrogenase are both physically and functionally coupled in mitochondrial complex II (Hancock et al., 2016). None of the other mitochondrial proteins investigated were found to be affected in the mutants, including UCP1, COXIV, Rieske FeS, NDUFA10 and beta F1 ATPase (Figures S6A to D). Levels of POX and UCP1 were not significantly changed by fasting in BAT of wild type mice (Figures S6E).

**Figure 5.**
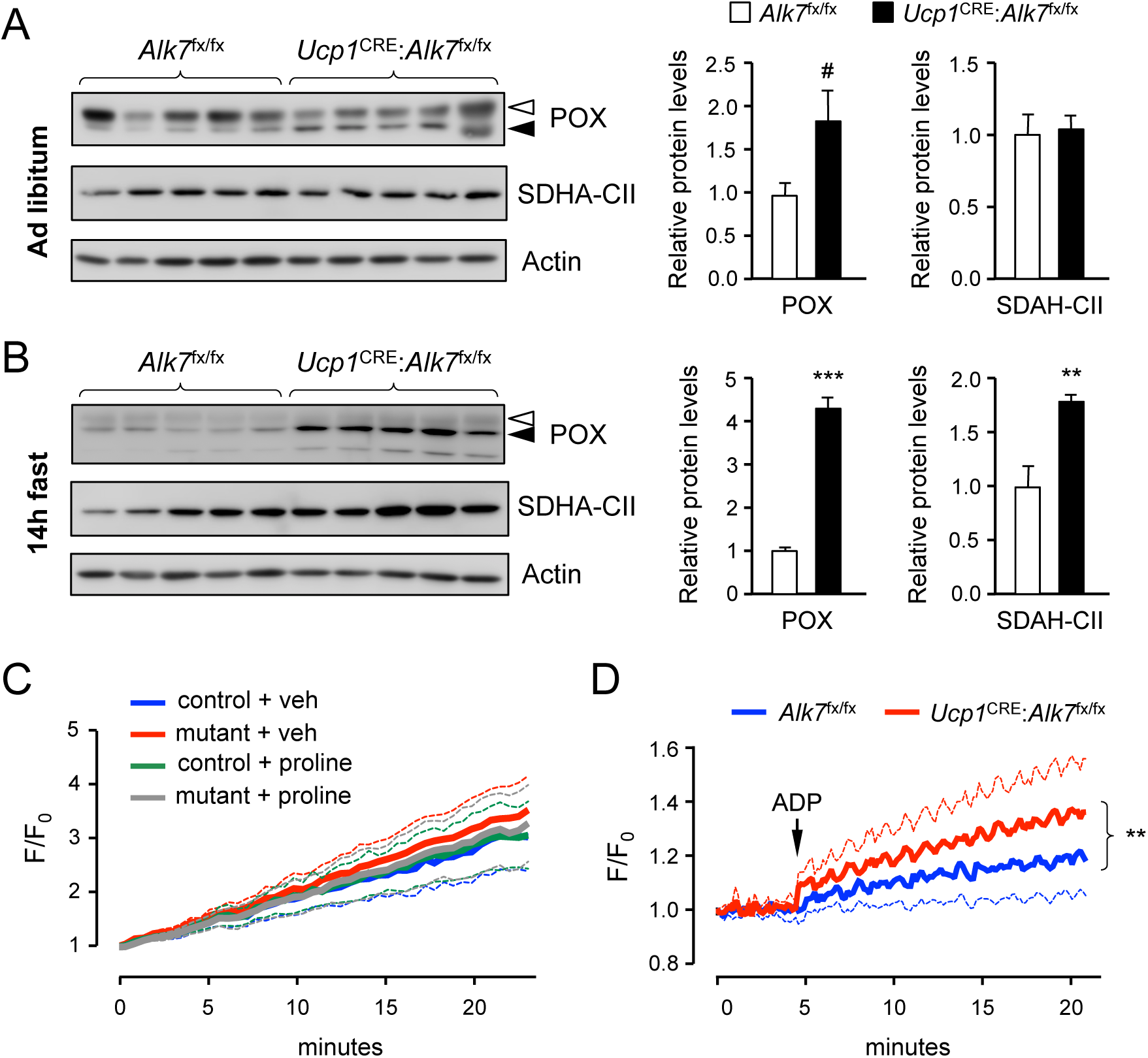
Increased proline-dependent ATP generation in mitochondria from iBAT lacking ALK7. (A, B) Western blot analysis of proline dehydrogenase (POX) and succinate dehydrogenase CII subunit (SDHA-CII) in iBAT of 2 month old conditional mutant and control mice fed Chow *ad libitum* (A) or after 14h fasting (B). Solid arrowheads point to POX band, open arrowheads denote unspecific band. Histograms to the right show quantitative analyses of protein levels normalized to those in control *Alk7*^fx/fx^ mice. #, P=0.069; **, P<0.01; ***, P<0.001; two-tailed unpaired Student’s t-test. (C) Traces of ROS production in mitochondria isolated from iBAT of 14h-fasted conditional mutant (red and grey) and control (blue and green) mice after vehicle (blue and red) or proline stimulation (green and grey). Dotted lines represent standard error. N=3 independent experiments. (D) Traces of proline-induced ATP production in mitochondria isolated from iBAT of 14h-fasted conditional mutant (red) and control (blue) mice. Dotted lines represent standard error. N=3 independent experiments. **, P<0.01; two-way ANOVA.

We investigated possible functional consequences of the increased levels of POX in BAT of fasted *Ucp1*^CRE^:*Alk7*^fx/fx^ mutant mice by assessing two of the most important effects attributed to proline oxidation by POX, namely production of reactive oxygen species (ROS) (Zarse:2012gh; Donald et al., 2001; Goncalves et al., 2014) and generation of ATP {Pandhare:2009ds; Phang:2017hs}. In control experiments, rotenone, a well known inducer of ROS production, produced a robust increase in ROS levels in BAT mitochondria (Figure S7A). However, in our hands, addition of proline to mitochondria isolated from iBAT of fasted mice failed to induce ROS production, regardless of genotype (Figure 5C). Next, we assessed ATP synthesis in BAT mitochondrial fractions from 14h fasted mutant and control mice. Compared to its effects on liver mitochondria (Figure S7B), proline supported a modest increase in ATP production in mitochondria isolated from BAT of fasted control mice (Figure 5D). However, ATP generation was significantly elevated in BAT mitochondria of fasted *Ucp1*^CRE^:*Alk7*^fx/fx^ mutant mice compared to controls (Figure 5D). These results suggest that nutrient stress induces elevated levels of POX which in turn lead to enhanced ATP generation in mitochondria from BAT lacking ALK7.

### Fasting-induced hypothermia in mice lacking ALK7 in brown adipocytes

In order to test the possible physiological significance of the changes observed in the metabolic functions of mutant iBAT lacking ALK7 we exposed control and mutant animals to acute cold (5°C for 4h) in metabolic cages and assessed energy consumption, body temperature and activity during this period. A second group of mice was fasted for 14h prior to cold exposure to test the effects of more stringent nutritional conditions. The body weights of mice placed in metabolic chambers was not different between genotypes. In fed mice, a 4h cold exposure did not reveal any alterations in either VO_2_, body temperature, RER (Figures 6A to C) or activity (Figure S8A) in the mutant mice compared to controls. The VO_2_ traces showed an expected increase during the first hour of exposure to cold, followed by a plateau that was largely maintained in the two genotypes (Figure 6A). However, significant changes were observed in animals that had been fasted for 14h prior to cold exposure. Mutant animals that had been fasted were unable to keep the higher levels of VO_2_ observed in control animals (Figure 6D) and, as a consequence, displayed a very rapid drop in body temperature (Figure 6E), with over 70% of animals showing temperatures lower than 32°C after 3h (compared to 20% of controls), a temperature at which rodents enter a state of torpor. Overall activity was similar to controls in the mutant mice during this period (Figure S8B). In these conditions, both genotypes showed a similar switch to usage of free-fatty acids (FFA) as energy source, as shown by RER close to 0.7 (Figure 6F), in line with the low levels of serum glucose and insulin observed upon cold exposure, which did not show differences between genotypes (Figures S8C and D). Both mutants and controls also showed similar serum levels of FFA and triglycerides (Figures S8E and F), indicating normal uptake of circulating lipids in the mutants (abnormal uptake is known to result in elevated levels in serum). We note that mutant mice exposed to 5°C for 8h with unrestricted access to food showed no difference in body temperature compared to controls (Figure S8G), indicating that the deficit in the mutants is specific to situations of nutritional stress. Lastly, we tested whether responses to norepinephrine (NE), a key activator of BAT upon cold exposure, were affected in the conditional mutants lacking ALK7 in brown adipocytes. NE content and turnover was comparable in iBAT from fasted mutant and control mice during cold exposure (Figure S9A). Also, *Ucp1* mRNA was similarly induced by cold exposure in both genotypes (Figure S9B), and levels of activated mTORC2 (phosphorylated at Ser^2481^) and HSL (phosphorylated at Ser^563^), which are targets of NE signaling in BAT (Albert et al., 2016), were similar in fasted mutant and control mice upon cold exposure (Figures S9C and D), suggesting normal NE signaling in mutant BAT. Together, these results suggest that hypothermia in mice lacking ALK7 in BAT may be the consequence of abnormally high catabolism of fat and amino acids upon nutrient stress, resulting in depletion of energy depots necessary to defend body temperature upon cold exposure.

**Figure 6.**
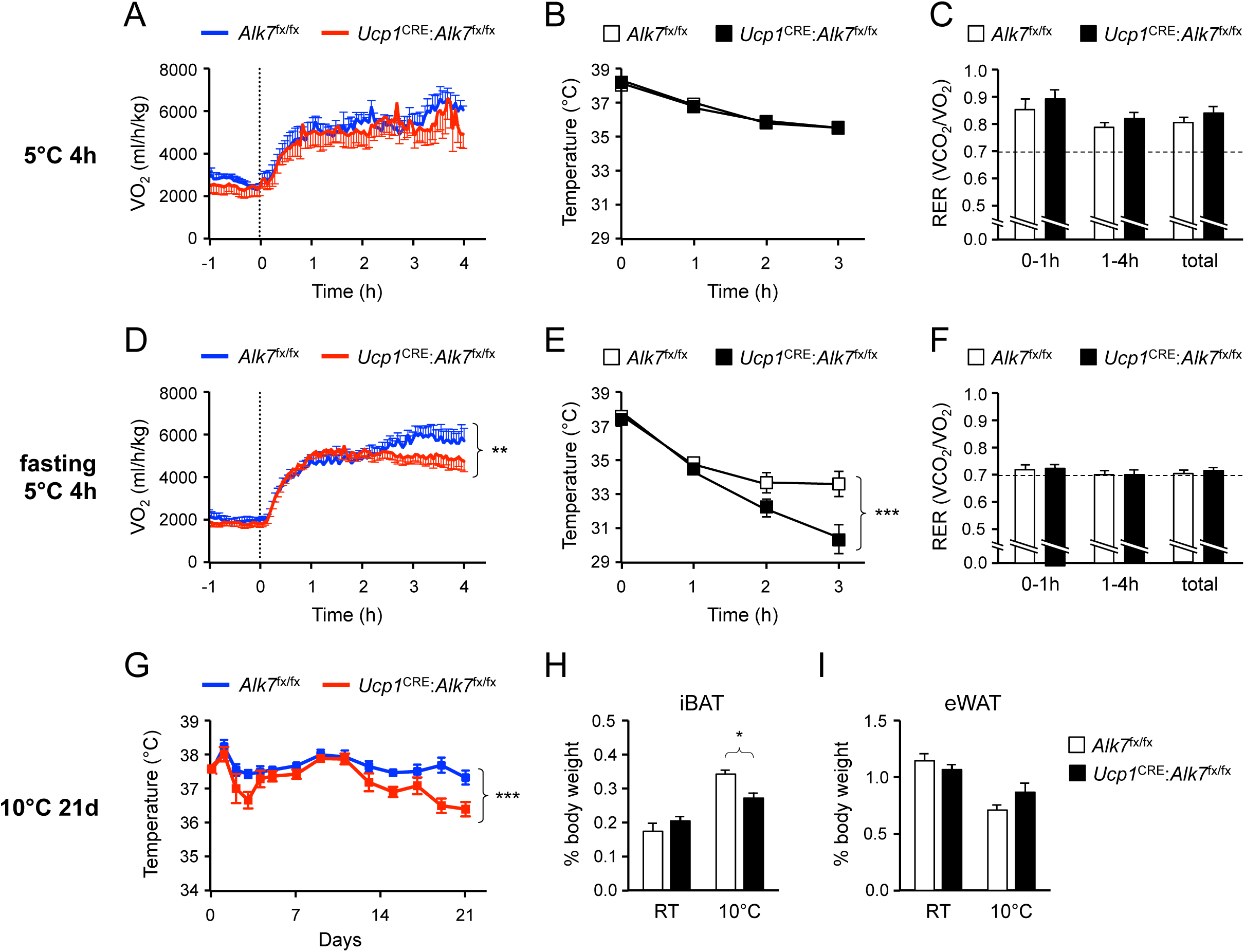
Fasting-induced hypothermia in mice lacking ALK7 in brown adipocytes. (A, D) VO_2_ measured by indirect calorimetry in *ad libitum* fed (A) or 14h-fasted (D) 2 month old conditional mutant and control mice during exposure to 5°C in metabolic cages. N=5 (A) and 10-12 (D) mice per group. **, P<0.01; two-way ANOVA (B, E) Rectal temperature in *ad libitum* fed (B) or 14h-fasted (D) 2 month old conditional mutant and control mice during exposure to 5°C. N=8 (B) and 10 (E) mice per group. ***, P<0.001; two-way ANOVA. (C, F) Respiratory exchange ratio (RER) measured by indirect calorimetry in *ad libitum* fed (C) or 14h-fasted (F) 2 month old conditional mutant and control mice during exposure to 5°C in metabolic cages. N=5 (C) and 10-12 (F) mice per group. (G) Rectal temperature in conditional mutant and control mice during chronic exposure to 10°C for 21 days (preceded by 14d acclimatization at 18°C). Temperature was measured every 2 days during the light phase of the day cycle. N=5 mice per group. ***, P<0.001; two-way ANOVA. (H, I) Relative iBAT and eWAT mass expressed as percentage of body weight in conditional mutant and control mice after 21d chronic cold exposure at 10°C. Values are presented as average ± SEM. N=5 mice per group. *, P<0.05; one-way ANOVA with Tukey’s post test.

In the final set of studies, we asked whether the metabolic defects underlying the inability of the mutants to maintain body temperature upon acute cold exposure in conditions of nutrient stress, could lead to impaired cold adaptation upon prolonged exposure to lower temperatures, even with unrestricted access to food. Control and mutant animals of matched age and weights were first cold adapted for 14 days at 18°C, and then placed at 10°C for 21 days. While control animals maintained body temperature throughout the experiment, a significant drop in body temperature was observed in the mutants during the last week of the treatment (Figure 6G). This was accompanied by a decrease in relative iBAT weight in the mutants exposed to cold, similar to that observed upon fasting, but not in eWAT weight (Figures 6H and I). Body weight and food intake were comparable in the two genotypes after cold exposure (Figures S10A and B). Also *Ucp1* mRNA was induced to similar levels by prolonged cold treatment in BAT of control and mutant mice (Figure S10C). Chronic cold exposure also induces *Ucp1* mRNA expression in subcutaneous WAT depots, particularly iWAT, a processed generally known as “browning”. We therefore examined expression of several thermogenic markers in iWAT of control and *Ucp1*^CRE^:*Alk7*^fx/fx^ mutant mice after prolonged cold exposure, but found that they were induced to comparable levels in both genotypes (Figures S11A to E), indicating normal browning of iWAT in conditional mutant mice after cold exposure. These results indicate that *Ucp1*^CRE^:*Alk7*^fx/fx^ mutant mice are inadequately adapted to prolonged cold exposure, perhaps due to premature depletion of energy depots in BAT, even in conditions of unrestricted food access.

## Discussion

Fasting triggers a range of catabolic activities that enable the usage of endogenous energy reservoirs, such as fat and proteins, allowing tissues to cope with their metabolic needs, ultimately contributing to the survival of the organism. Unlike liver, muscle and other tissues, our understanding of the physiological signals that adapt BAT function to different nutritional states is very limited. Based on the results of the present study, we propose that ALK7 functions to dampen catabolic activities triggered in BAT upon limited nutrient availability. Under nutrient stress, these catabolic functions, e.g. lipolysis and amino acid degradation, become abnormally amplified in brown adipocytes lacking ALK7, leaving the tissue energetically unable to cope with the demands imposed by low ambient temperatures. Recent studies have reported that circulating energy substrates are sufficient to fuel non-shivering thermogenesis under conditions that blunt BAT lipolysis, even upon acute cold exposure in the absence of food (Schreiber et al., 2017; Shin et al., 2017), leading to the notion that lipid droplet lipolysis in brown adipocytes is not essential for cold-induced thermogenesis regardless of food availability (Shin et al., 2017). However, these studies left open the question of the importance of endogenous BAT stores when animals confront lower temperatures after a previous period of prolonged fasting. Under such more stringent nutrient conditions, which we presume not to be uncommon in the wild, our findings indicate that energy reservoirs within BAT become crucially important to maintain body temperature. Our study indicates that an intrinsic abnormality in the catabolic function of BAT can indeed compromise responses to cold exposure.

Fasting induces expression of *Klf15* in liver, where it upregulates gluconeogenesis, and in muscle, where it promotes amino acid degradation, thereby providing precursors for liver gluconeogenesis at the expense of muscle protein (Yamamoto et al., 2004; Gray et al., 2007; Shimizu et al., 2011). Glucocorticoid signaling appears to be responsible for *Klf15* induction upon fasting in these tissues (Gray et al., 2007; Sasse et al., 2013). In contrast, *Klf15* expression was recently reported to be decreased in white adipocytes from fasted mice compared to fed mice (Matoba et al., 2017). The same study showed that WAT KLF15 inhibits lipolysis and promotes lipid storage in response to insulin, indicative of anabolic functions in this tissue. On the other hand, the regulation and possible functions of KLF15 in BAT have been unknown. Unlike WAT, and in line with the observations made in liver and muscle, we find that fasting induces *Klf15* gene expression in BAT, which, suppression by RU486, suggests it to also be under the control of glucocorticoid signaling. Interestingly, muscle and BAT have been shown to originate from a common set of dermomyotome-derived precursor cells, which are distinct from those that give rise to WAT (Wang and Seale, 2016), suggesting a possible reason for their sharing a similar mode of *Klf15* regulation upon fasting. Our study reveals a novel signaling pathway of ALK7 function in BAT involving glucocorticoid, KL15 and POX, which is distinct from its known role in regulating adrenergic activity in WAT (Guo et al., 2014).

Previous studies in cancer cells have shown that POX can promote cell survival under conditions of limited nutrient availability through its ability to catabolize collagen-derived proline (Olivares et al., 2017). Proline is the most abundant amino acid in collagen, itself a major component of the extracellular matrix of many tissues, including BAT. Under conditions of low fatty acid and glucose availability, increased POX expression in BAT lacking ALK7 may lead to channeling of TCA intermediates and mitochondrial generation of ATP, as observed in fasted mutant mice, leaving these animals with fewer reserves to successfully confront exposure to cold temperatures. In line with this, the body temperature of fasted mutant mice dropped significantly after 3h cold exposure, in parallel with lower energy expenditure.

Mutant mice lacking ALK7 in BAT were unable to maintain normal BAT mass and body temperature under chronic cold exposure (3 weeks), despite unrestricted access to food. We note that patients undergoing chronic (2 weeks) glucocorticoid treatment showed reduced BAT mass compared to controls (Ramage et al., 2016). It is possible that chronic exposure to overactive glucocorticoid-regulated pathways in BAT of conditional mutant mice may exaggerate BAT catabolic functions, resulting in reduced BAT mass and lower thermogenic performance, even under normal feeding conditions. At any rate, these observations indicate an important role for BAT ALK7 in the adaptation to chronic cold exposure.

In summary, we have discovered a novel role for the TGF-β superfamily receptor ALK7 in the adaptation of BAT physiology to variations in nutritional status, to our knowledge, the first mechanism described to regulate this important process. We find that ALK7 functions by limiting nutrient stress-induced overactivation of catabolic activities in brown adipocytes. Mechanistically, it does so by suppressing the levels of ATGL, a key enzyme for lipolysis, as well as a novel pathway involving downstream effectors of glucocorticoid signaling, such as KLF15, a master regulator of fasting responses, and POX, another nutrient-sensitive component of survival responses to fasting. A better understanding of the mechanisms by which BAT adapts to fluctuations in nutrient availability will be critical for the development of safe methods to harness energy expenditure in BAT to combat human obesity and metabolic syndrome.

## Materials and Methods

### Animals

Mice were housed under a 12h light-dark cycle, and fed a standard chow diet or a high-fat diet (HFD, 60% of calorie from fat; ResearchDiet D12492). The mouse lines utilized in this study have been described previously and are as follows: i) conditional *Alk7*^fx/fx^ (Guo et al., 2014); ii) global knock-out *Alk7*^-/-^ (Jörnvall et al., 2004); and iii) CRE line *Ucp1*^CRE^ (Kong et al., 2014); all back-crossed for at least 10 generations to a C57BL/6J background (considered as wild type). Animal protocols were approved by Stockholms Norra Djurförsöksetiska Nämnd (Stockholm North Ethical Committee for Animal Research) and are in accordance with the ethical guidelines of the Karolinska Institute.

### Cold exposure, calorimetry and body composition

For cold exposure, animals were housed in a Memmert HPP750 climate chamber at the indicated temperatures. For acute cold exposure, body temperature was measured every hour using a rectal thermometer. After 3h at 5°C, animals were sacrificed and iBAT extracted for molecular studies. For chronic cold exposure, animals were acclimatized to 18°C for 14 days prior to 21d exposure to 10°C. Body temperature was measured every 2 days during the light phase of the day cycle. Indirect calorimetry, food intake, and locomotor activity were assessed with a PhenoMaster Automatic Home Cage system (TSE Systems). Mice were housed individually with ad libitum access to food and water. Mice were acclimatized to the metabolic cages prior to automated recordings. For acute cold exposure in metabolic cages, mice were placed 4h at 5°C. Fat and lean mass were measured using a body composition analyzer EchoMRI-100TM.

### Insulin sensitivity, RU486 treatment and NE turnover

For insulin sensitivity test, mice fasted for 4 hours received an intraperitoneal injection of 0.75 U/kg Humulin-R insulin (Eli Lilly) or saline (vehicle). Mice were sacrificed 1h later, and tissue samples were snap-frozen at -80 °C for subsequent analysis. Ru486 (Sigma-Aldrich) was freshly formulated in DMSO before use. Weight-paired mice were fasted for 10h and then injected intraperitoneally with 5mg of RU486 in 50μl of DMSO or vehicle (only DMSO). Mice were sacrificed 4h later, and tissue samples were snap-frozen at -80 °C for subsequent analysis. For analysis of NE turnover, mice fasted overnight (14h) were injected with 250mg/kg of the norepinephrine synthesis blocker α-Methyl-DL-tyrosine methyl ester hydrochloride (AMT, Sigma-Aldrich) and immediately placed at 5°C. Mice were sacrificed after 3h cold exposure and tissue samples were snap-frozen at -80 °C for subsequent NE quantification. Tissue NE was measured by ELISA kit (Labor Diagnostika Nord) according to the manufacturer’s recommendation.

### Measurements in blood and serum samples

Blood samples were obtained by tail tip bleeding in the morning. Blood glucose was determined using a glucometer (Accutrend; Roche). Serum from blood was obtained by centrifugation of blood at 9,391xg for 15min at 4°C. The pellet was discarded and serum samples were stored at -80° C for further analysis. Free fatty acid and triglyceride levels were measured in serum using the colorimetric quantification kits Half-micro test (Roche) and Infinity (ThermoScientific), respectively. Insulin was measured in serum with an ultra-sensitive mouse insulin ELISA kit (Mercodia). Corticosterone in serum was measured by ELISA kit (Enzo). All commercial kits were used following the manufacturer’s recommendations.

### Ex vivo lipolysis

For ex vivo lipolysis, iBAT was dissected from mice that were either fed ad libitum or fasted for 6h, and tissue pieces were placed in DMEM at 37°C for 1h. The medium was then replaced with DMEM containing 2% fatty-acid-free BSA for 1h at 37°C. After incubation, the medium was collected and iBAT pieces were solubilized in 0.3N NaOH/0.1% SDS at 65°C overnight and subsequently centrifuged at 845xg for 15min at 4°C to remove the layer of fat. Protein content was determined using Pierce BCA Protein assay (Pierce, Thermo Fisher Scientific). Glycerol release to the media was measured using a free glycerol reagent (Sigma-Aldrich), and normalized to the total amount of protein in the tissue samples.

### Histological analysis of iBAT sections

iBAT tissue samples were fixed in 4% PFA, embedded in 4% agar and cut into 50 µm-thick sections in a vibratome. Sections were incubated in 10% sucrose for 15min, followed by 30% sucrose for 3h at 4 °C, and permeabilization by 4 consecutive freeze-thaw cycles in 30% sucrose. Sections were stained with 10µg/ml BODIPY 493/503 and mounted onto glass slides for imaging a confocal microscope (Zeiss). The area individual lipid droplets was measured with Zen software (Zeiss) and used for quantification of lipid droplet size.

### Isolation of mitochondria and measurements of ATP synthesis and ROS production

For mitochondria isolation, iBAT and liver were dissected out, washed and minced in BAT (250mM sucrose and 0.1% fatty-acid-free BSA) or liver (125mM sucrose, and 0.1% fatty-acid-free BSA) isolation buffer, respectively. The tissue was homogenized in a glass homogenizer in isolation buffer supplemented with protease inhibitors (Roche). Homogenates were filtered through a 70μm mesh and nuclei were removed by centrifugation at 845xg for 10min at 4°C in a microcentrifuge. Mitochondrial fractions were then collected by centrifugation at 9,391xg for 15min at 4°C, resuspended in MSK buffer (75mM mannitol, 25mM sucrose, 5mM potassium phosphate, 20mM Tris-HCl, 0.5mM EDTA, 100mM KCl, and 0.5% fatty-acid-free BSA, pH 7.4) and kept on ice until used. Proteins were measured by the BCA method.

ATP synthesis was determined fluorometrically in isolated mitochondria using a coupled enzyme assay with continuous monitoring of the reduction of NADP as described previously (Korge et al., 2003), with minor modifications. Mitochondrial fractions (0.5mg/ml) were suspended in 150µl of MSK buffer in the presence of 5mM proline, 1mM NADP, 10mM glucose, 10U/ml hexokinase, 5U/ml glucose-6-P dehydrogenase, and 0.5% fatty-acid-free BSA. ATP synthesis was started by the addition of 100μM ADP and measured as an increase in NADPH fluorescence (excitation=340 nm, emission=450 nm) at 37°C under constant stirring in a SpectraMax M2 microplate reader (Molecular Devices).

Mitochondrial ROS was measured as described previously with slight modifications (Goncalves et al., 2014). Mitochondrial fractions (0.01mg/ml) were suspended in 0.15ml of MSK buffer supplemented with 0.5% fatty-acid-free BSA in the presence of 12U/ml horseradish peroxidase, 45U/ml superoxide dismutase, and 50µM Amplex UltraRed. Superoxide production was started by the addition of 5mM proline and converted to H_2_O_2_ by superoxide dismutase. Appearance of H_2_O_2_ was monitored as the increase in fluorescence of the oxidized form of Amplex UltraRed (excitation=545 nm, emission=600 nm) at 37°C under constant stirring in a SpectraMax M2 microplate reader (Molecular Devices).

### Isolation and mass spectrometry analysis of lipid droplets

Lipid droplets were isolated and delipidated as described previously (Brasaemle and Wolins, 2016) with slight modifications. Tissue was placed in ice-cold hypotonic lysis medium (HLM, 10 mM Tris, pH 7.4, 1 mM EDTA) in the presence of a protease inhibitor cocktail (Roche), minced and homogenized by 20 strokes in a glass homogenizer. Lysates (≈11 ml) were centrifuged at 26,000xg for 30 min at 4°C in a SW41Ti rotor (Beckman). The floating lipid droplet layers were harvested with a glass pipette and adjusted to 25% sucrose and 100 mM sodium carbonate, pH 11.5, using 60% sucrose and 1M sodium carbonate stock solutions containing protease inhibitors, by gentle mixing by pipetting. These fractions (≈4 ml) were layered into centrifuge tubes containing 1ml cushions of 60% sucrose and then overlaid with ≈5ml of 100mM sodium carbonate, pH 11.5, with protease inhibitors followed by ≈1 ml of hypotonic lysis medium with protease inhibitors. Tubes were centrifuged at 26,000xg for 30min at 4°C in a SW41Ti rotor. Floating lipid droplets were harvested using a pipette tip into 2-ml microcentrifuge tubes. Residual carbonate solution was removed by centrifuging tubes at 11,363xg for 20min at 4°C in a microcentrifuge; the lower fraction was removed with an 18-gauge needle from below the floating lipid droplet fraction. Lipid droplet fractions in microcentrifuge tubes were delipidated with 2 l of cold acetone overnight at -80 °C, followed by centrifugation at 11,363xg for 30min at 4°C, and removal of solvent from the protein pellet. The pellet was further extracted with acetone at room temperature, followed by 1:1 acetone:ether (v:v), and finally ether. Residual solvents were allowed to evaporate completely and lipid droplet fractions were stored at -80 °C. Mass spectrometry was performed on 4 separate sample sets, each set consisting of lipid droplets isolated from iBAT pooled from 2-3 age-paired *Alk7*^-/-^ and wild type mice. Delipidated lipid droplet pellets were resuspended using urea, sonication and vortexing. A tryptic digestion of 10µg was carried out with Urea/proteaseMax protocol for subsequent nLC-MS/MS analysis on QExactive, long gradient and nLC II. A standard proteome quantitation analysis was then performed.

### Primary culture of mouse and human brown adipocytes

iBAT from 4-6 month old mice was dissected out, micd and digested for 60min in DMEM/F12 medium supplemented with 1% BSA, antibiotics, and 1mg/ml collagenase II (Sigma Aldrich) under constant shaking. The digested tissue was filtered through 250µm nylon mesh and a 70µM cell strainer and centrifuged for 10min at 1500rpm to separate floating mature adipocytes. The pellet was resuspended in erythrocyte lysis buffer (15 mM NH4Cl, 10mM KHCO3, 0.1mM EDTA) for 10min to remove blood cells. The cells were further centrifuged at 1500rpm for 10 min, and the pellet (stromal vascular fraction, SVF) was resuspended and plated in 48-well plates. Cells were grown in DMEM/F12 supplemented with 10% FBS and 100µg/ml penicillin-streptomycin at 37°C until confluence. Adipocyte differentiation was induced (day 0) in medium supplemented with 1µM dexamethasone, 66nM insulin, 15mM HEPES, 1nM T3, 33µM biotin, 17µM pantothenate, 10µg/ml transferrin, and 1µg/ml rosiglitazone until full differentiation (day 10).

Human brown preadipocytes (kindly provided by Dr. Shingo Kajimura) were differentiated as previously described (Shinoda et al., 2015). Cells were grown in maintenance medium (Advanced DMEM/F12 supplemented with 10% FBS, 100µg/ml penicillin-streptomycin, 1nM T3, 2µg/ml dexamethasone and 5µg/ml insulin) at 37°C until confluence. Differentiation was started (day 0) with induction medium (Advanced DMEM/F12 supplemented with 10% FBS, 100µg/ml penicillin-streptomycin, 1nM T3, 2µg/ml dexamethasone, 5µg/ml insulin, 1µM rosiglitazone, 0.125mM indomethacin, and 0.25mM IBMX) in collagen-coated 48-well plates until full differentiation (day 24).

For activin B treatment, fully differentiated mouse or human brown adipocytes were incubated with 100ng/ml activin B (R&D Systems) in DMEM/F12 supplemented with 10% FBS and 100µg/ml penicillin-streptomycin for 24h before lysis and for RNA extraction of RNA (as indicated below). SB-431542 (Sigma-Aldrich) was added at the same time at a final concentration of 10µM, as indicated.

### RNA isolation, real-time quantitative PCR and microarray analysis

RNA from tissue and cells was extracted using the RNeasy Mini Kit (Qiagen), treated with DNase I (Life Technologies), and reversely transcribed using SuperScript II reverse transcriptase (Life Technologies). cDNAs were used for real-time quantitative PCR (Q-PCR) using the primers listed in Supplementary Table 1. Q-PCR was performed with SYBR Green PCR Master Mix (Life technologies) on a StepOnePlus real time PCR system (Applied Biosystems), using 18S rRNA as an endogenous normalization control. For quantification of mitochondria DNA, total tissue DNA was extracted with a DNeasy kit and used for real-time quantitative PCR using primers for the mitochondrial gene *CytB*, encoding cytochrome B (Guo et al., 2014).

### Western blotting

Snap-frozen tissues were homogenized with a high-speed tissue homogenizer in ice-cold RIPA buffer (50mM Tris-HCl, 150mM NaCl, 5mM EGTA, 1% NP-40, pH7.4) supplemented with proteinase and phosphatase inhibitor cocktails (Roche), and centrifuged at 845xg for 15min at 4°C to collect supernatants. Protein concentration was determined by the BCA method. Supernatants were used for reducing SDS-PAGE, transferred onto PVDF membranes (Amersham) and analyzed by Western blotting using the specific first antibodies listed in Supplementary Table 2 with goat anti-rabbit and goat anti-mouse peroxidase-conjugated immunoglobulin as secondary antibodies (Dako). The blots were then processed with the luminescence technique ECL (Thermo Scientific), and imaged in an Imagequant LAS4000. The levels of target proteins were quantified by the intensity of Western blot bands using ImageJ software (National Institutes of Health), using actin as loading control.

### Statistical analyses

Statistics analyses were performed using Prism 5 software (GraphPad, SPSS IBM corporation) and Microsoft Excel (Microsoft). Student’s t test, one-way ANOVA or two-way ANOVA were performed to test statistical significance according the requirements of the experiment. In some cases, two-tailed Mann Whitney was used as a non-parametric test. Bonferroni or Tukey’s post-tests were used, when indicated, as a further test for experiments that required multiple comparisons. The level of statistical significance was set at P<0.05 for all the analyses (*). All P values are reported in the figure legends.

## Acknowledgements

We thank Annika Andersson (Karolinska Institute) for help with mice genotyping; Dorothea Rutishauser (Karolinska Institute) for help with mass spectrometry analysis of lipid droplet proteomes; Julie Massart and Marie Björnholm (Karolinska Institute) for help with metabolic cages; Boon Seng Soh, Kian Leong Lee, Henry Yang He, and Bing Lim (ASTAR, Singapore) for help with microarray studies; Jan Nedergaard and Barbara Cannon (Stockholm University) for advice and help with preliminary studies; and Evan Rosen (Harvard Medical School) for *Ucp1*^CRE^ mice. Support for this research was provided by grants to C.F.I. from the Swedish Research Council, Swedish Cancer Society, Knut and Alice Wallenbergs Foundation (Wallenberg Scholars Program), National Medical Research Council of Singapore and National University of Singapore.

## Author contributions

P.M.-C. performed all the experiments, except the microarray study, which was done by C.F.I. P.M.-C. and C.F.I. designed the experiments and performed data analysis. P.M.-C. prepared a draft of the manuscript and figures. C.F.I. wrote the final paper.

---

**Supplementary Figure 1.**
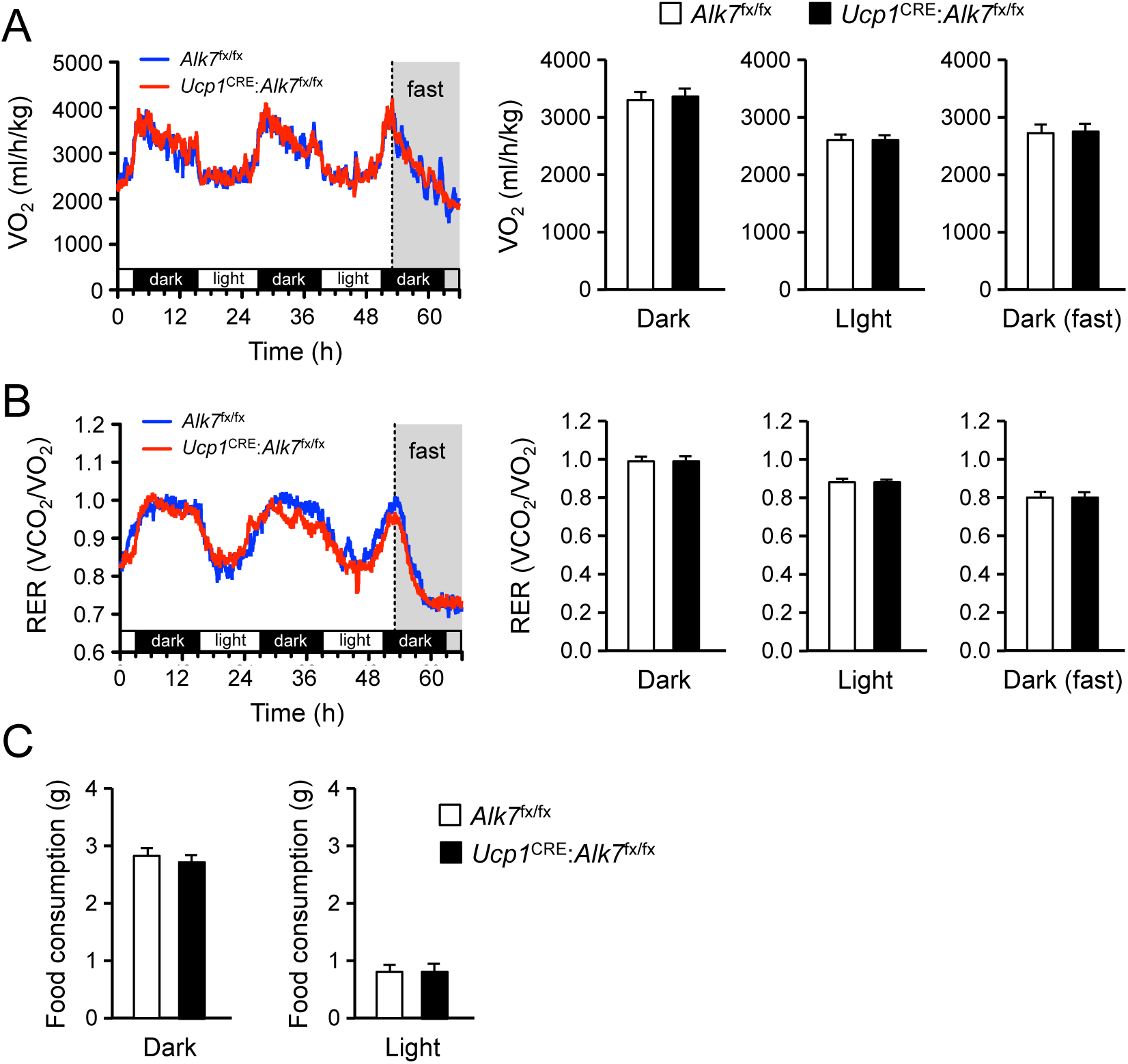
Normal energy consumption and food intake in mutant mice lacking ALK7 in BAT. (A, B) VO_2_ and RER measured by indirect calorimetry at room temperature in 2 month old mice. RER values close to unit indicate carbohydrate usage, while values around 0.7 indicate fat consumption. Animals were fasted during the last 14h of the experiment (N = 10-12 mice per group). (C) Food consumption in 12h dark or light cycle at room temperature N=14-16 mice per group.

**Supplementary Figure 2.**
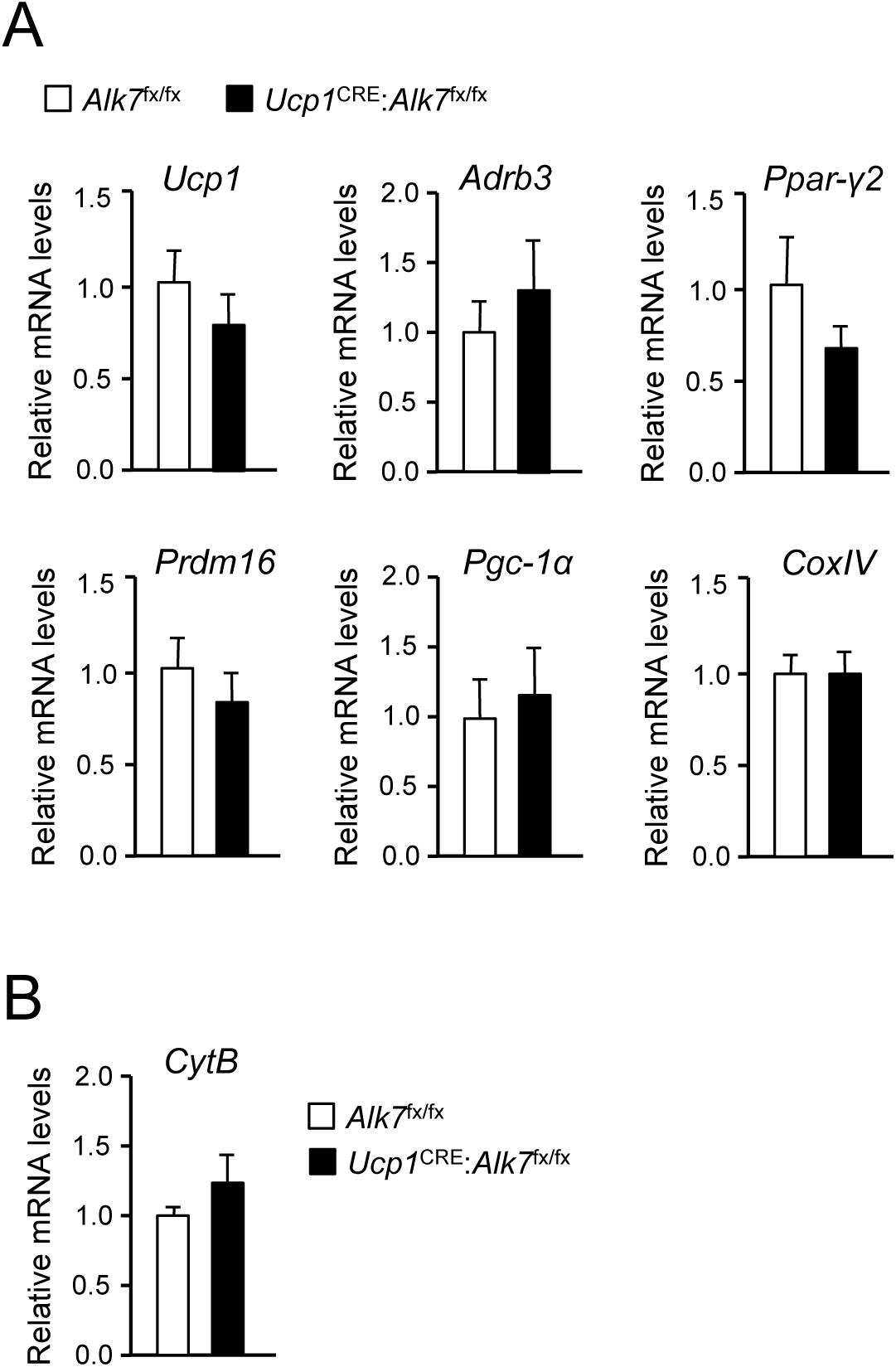
Normal BAT differentiation in mice lacking ALK7 in brown adipoctes. (A) mRNA expression of differentiation and maturation markers in BAT of 2 month old control (*Alk7f*^x/fx^) and mutant (*Ucp1*^CRE^:*Alk7*^fx/fx^) mice assessed by Q-PCR. Values are expressed relative to levels in control mice. *Ucp1*, Uncoupling protein-1; *Adrb3*, Adrenergic recetor beta 3; *Ppar-γ2*, Peroxisome proliferator-activated receptor gamma 2; *Prdm16*, PR/SET Domain 16; *Pgc-1α*, Peroxisome proliferator-activated receptor-gamma coactivator 1 alpha; *CoxIV*, Cytochrome C oxidase subunit 4. N=4-5 mice per group. (B) Mitochondrial DNA content was determined by quantitative PCR using primers specific for the mitochondrial gene *CytB*, Cytochrome B. N=5 mice per group.

**Supplementary Figure 3.**
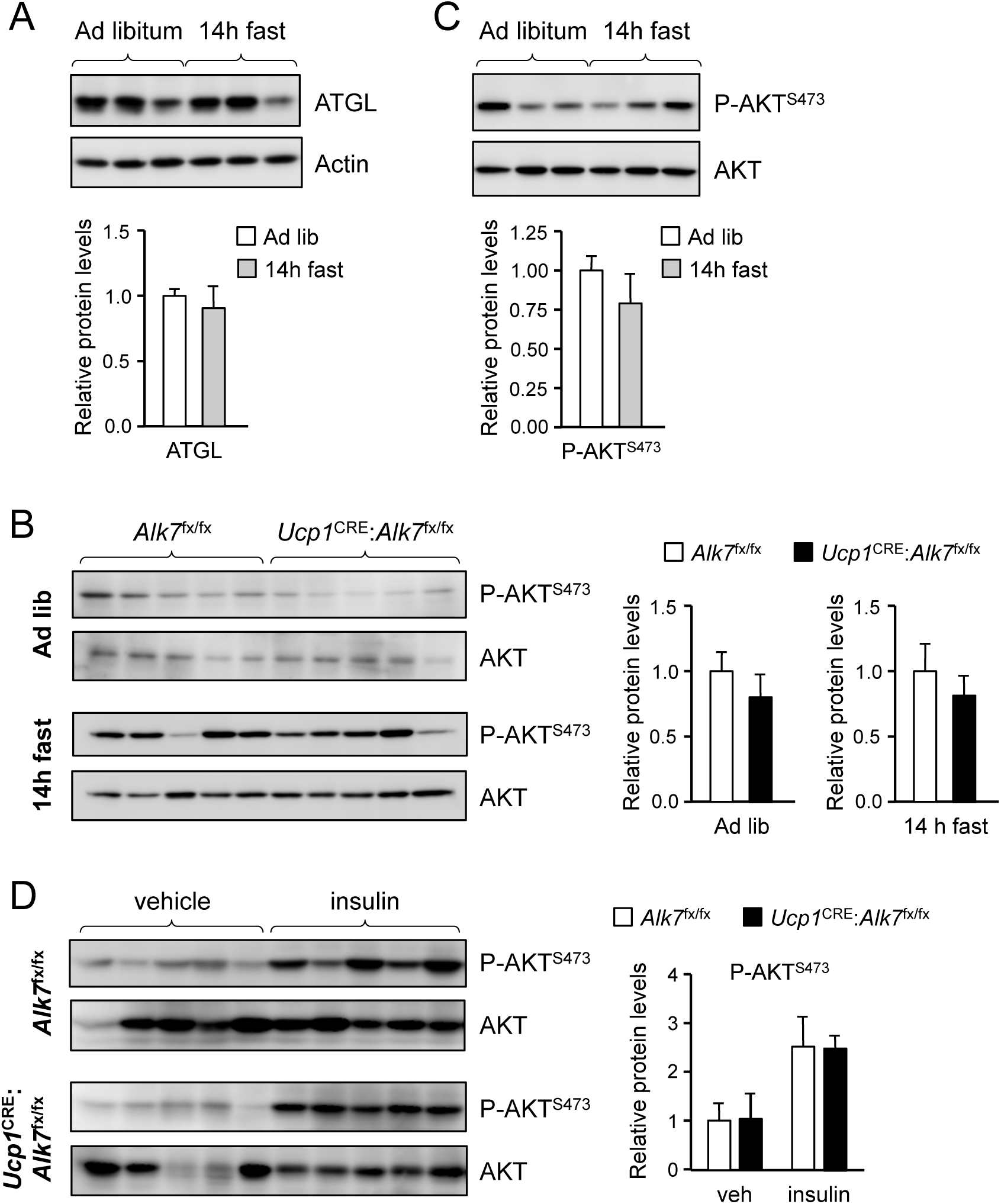
Unchanged levels of ATGL and P-AKT in wild type mice upon fasting and normal P-AKT levels and insulin sensitivity in iBAT lacking ALK7. (A) ATGL in BAT of wild type mice fed on Chow ad libitum or fasted for 14h, assessed by Western blotting of BAT lysates. Histogras shows ATGL normalized to actin relative to ad libitum. N=3 mice per group. (B) Phospho-AKT^S473^ in BAT of control (*Alk7*^fx/fx^) and mutant (*Ucp1*^CRE^:*Alk7*^fx/fx^) 2 month old mice fed on Chow ad libitum (Ad lib) or fasted for 14h, assessed by Western blotting of BAT lysates. Histograms show P-AKT^S473^ levels normalized to total AKT relative to control. N=5 mice per group. (C) Phospho-AKT^S473^ in BAT of wild type mice fed on Chow ad libitum or fasted for 14h, assessed by Western blotting of BAT lysates. Histogram shows P-AKT^S473^ normalized to total AKT relative to ad libitum. N=3 mice per group. (D) Phospho-AKT^S473^ in BAT of control (*Alk7*^fx/fx^) and mutant (*Ucp1*^CRE^:*Alk7*^fx/fx^) 2 month old mice following injection with insulin or vehicle assessed by Western blotting of BAT lysates. Histograms show P-AKT^S473^ levels normalized to total AKT relative to control. N=5 mice per group.

**Supplementary Figure 4.**
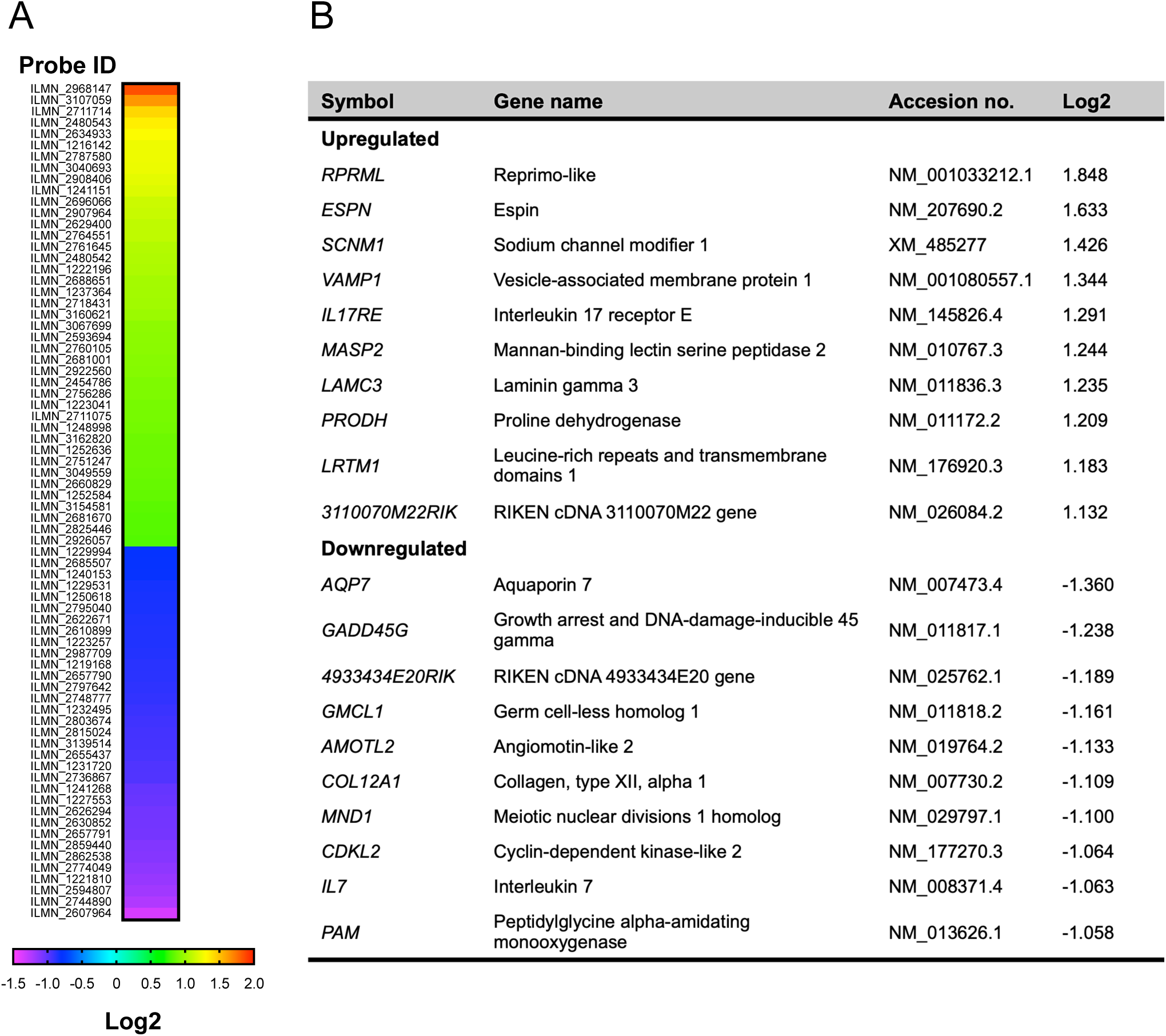
Microarray analysis of genes differentially expressed in iBAT of *Alk7*^-/-^ global knock-out mice compared to wild type. (A) Heat map showing relative expression of mRNAs differentially expressed (P<0.05) in iBAT of *Alk7^-/-^* global knock-out mice relative to wild type controls. The animals (N=4 per group) were kept at 30°C prior to iBAT extraction. Illumina microarray Probe IDs are indicated. Values are expressed as Log2 ofknock-out to wild type expression ratio. (B) Top 10 up- and down-regulated genes in iBAT of *Alk7^-/-^* mice compared to wild type.

**Supplementary Figure 5.**
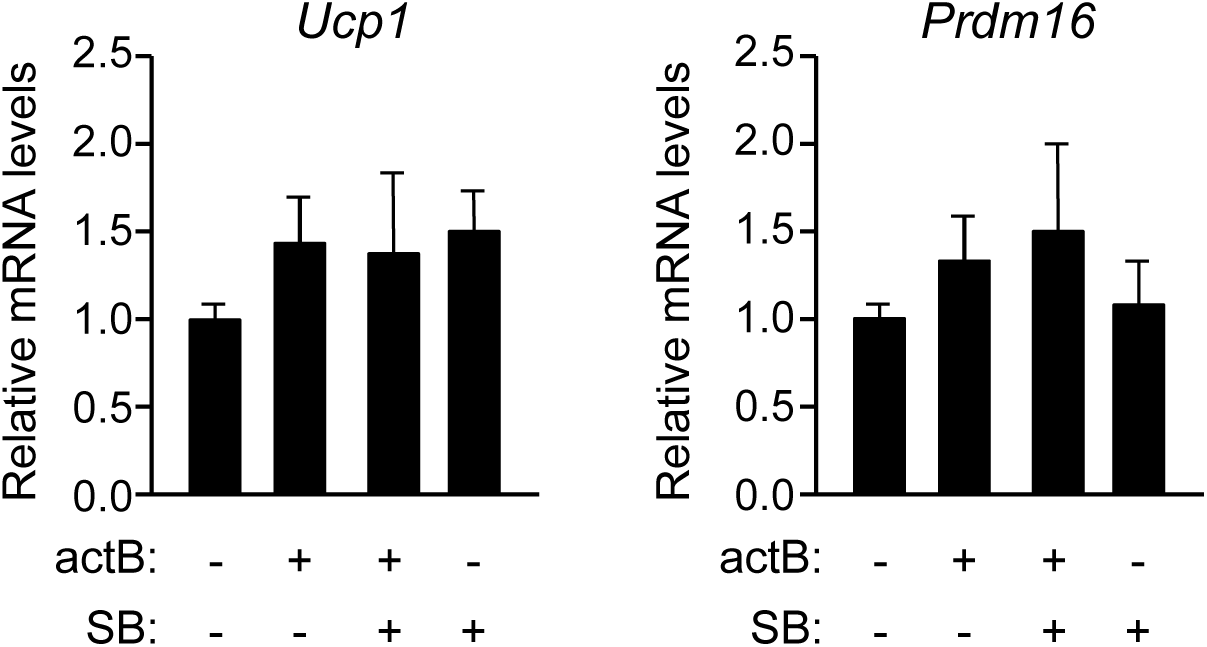
Effect of activin B on mRNA expression of BAT markers *Ucp1* and *Prdm16 assessed* in cultured brown adipocytes. Relative levels of *Ucp1* and *Prdm16* mRNAs were assessed by Q-PCR after 24hstimulation with activin B in the presence or absence of SB431542 inhibitor as indicated. Values were normalized to control untreated cutlures. Shown are average ± SEM of N=5 independent experiments each performed in duplicate.

**Supplementary Figure 6.**
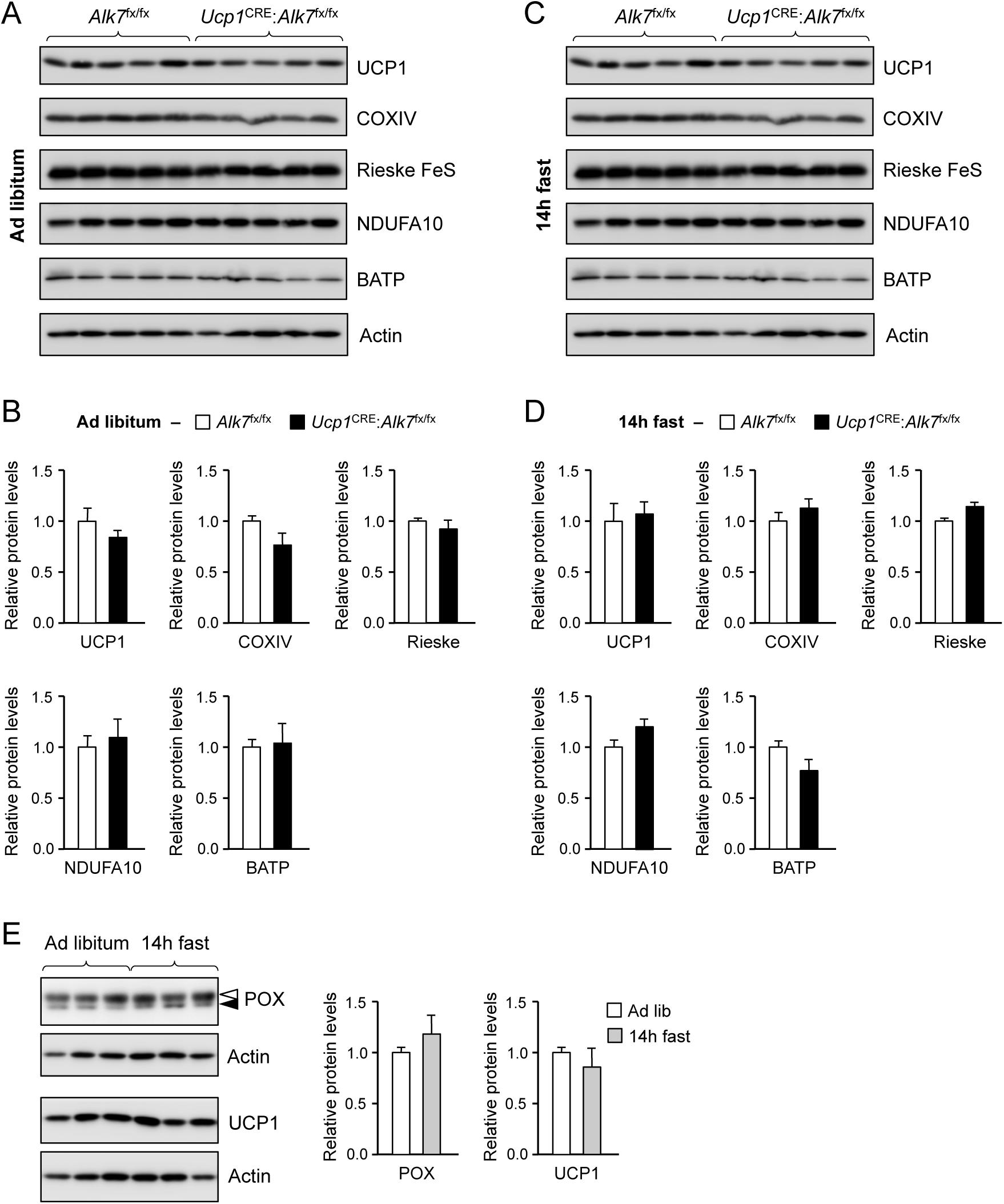
Expression of mitochondrial proteins in BAT of control and conditional mutant mice fed ad libitum or after 14h fasting. (A) Expression of UCP1, COXIV, Rieske FeS, NDUFA10 and beta F1 ATPase (BATP) in BAT of control (*Alk7*^fx/fx^) and mutant (*Ucp1*^CRE^:*Alk7*^fx/fx^) 2 month old mice fed on Chow ad libitum (Ad lib) assessed by Western blotting of BAT lysates. (B) Histograms showing levels of the indicated proteins normalized to actin relative to control in *Alk7*^fx/fx^ (white bars) and mutant (black bars) mice fed ad libitum. N=5 mice per group. (C) Expression of UCP1, COXIV, Rieske FeS, NDUFA10 and beta F1 ATPase (BATP) in BAT of control (*Alk7*^fx/fx^) and mutant (*Ucp1*^CRE^:*Alk7*^fx/fx^) 2 month old mice fasted for 14h assessed by Western blotting of BAT lysates. (D) Histograms showing levels of the indicated proteins normalized to actin relative to control in *Alk7*^fx/fx^ (white bars) and mutant (black bars) mice fasted for 14h. N=5 mice per group. (E) POXand UCP1 protein in BAT of wild type mice fed on Chow ad libitum or fasted for 14h, assessed by Western blotting. Solid arrowheads point to POX band, open arrowheads denote unspecific band. Histogram shows POX normalized to actin relative to ad libitum. N=3 mice per group.

**Supplementary Figure 7.**
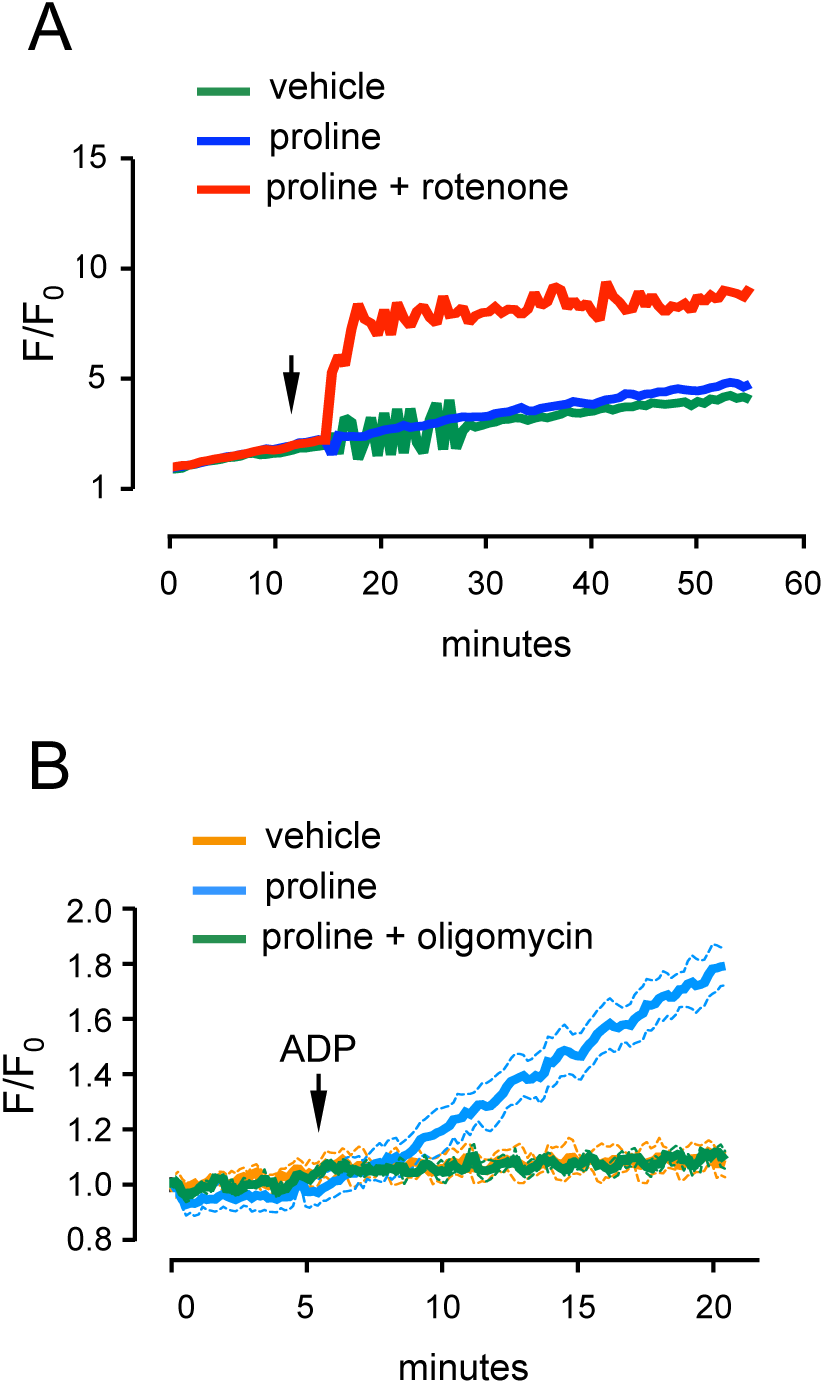
Control experiments for measruments of ROS and ATP production. (A) ROS production in BAT mitochondria from wild type mice induced by rotenone and proline (arrow). N=3 independent mitochondrial preparations. (B) ATP generation in liver mitochondria from fasted wild type mice was robustly induced by ADP (arrow) in the presence of proline (blue) and was blocked by addition of the ATP synthase inhibitor oligomycin (green). N=3 independent mitochondrial preparations.

**Supplementary Figure 8.**
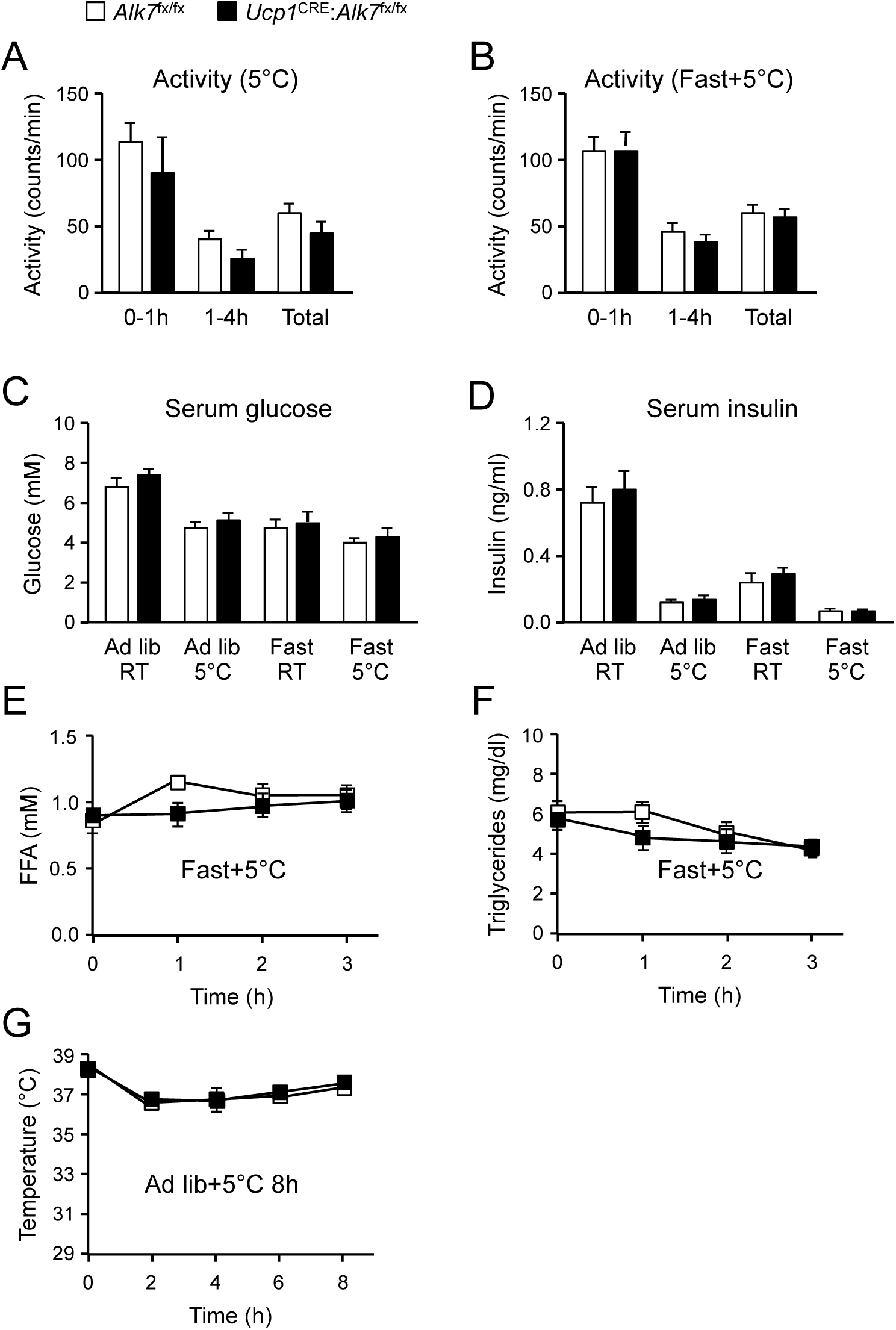
Influence of fasting and cold exposure on levels of glucose, insulin, activity, free-fatty acids, triglycerides and body temperature. (A, B) Activity in metabolic chambers of mice during exposure at 5°C preceded by *ad libitum* feeding (panel A, N=5 mice per genotype) or after 14h fasting (panel B, N=10 mice per genotpye). (C) Serum glucose levels under different feeding and temperature conditions. Fasting was 14h and cold exposure was 4h. Values show averages ± SEM. N=15, (ad lib RT); 5, ad lib 5°C; 10, fast RT; 10, fast 5°C. (D) Serum insulin levels under different feeding and temperature conditions. Fasting was 14h and cold exposure was 4h. Values show averages ± SEM. N numbers as above. (E, F) Serum free fatty acids (FFA, panel E) and triglycerides (F) in control and conditional mutant mice exposed to 5°C for 3h after 14h fasting. Shown are averages ± SEM. N=6 mice per group. (G) Rectal temperature in control and conditional mutant mice during 8h exposure at 5°C fed *ad libitum* both before and durign cold exposure. Shown are averages ± SEM. N=5 mice per group.

**Supplementary Figure 9.**
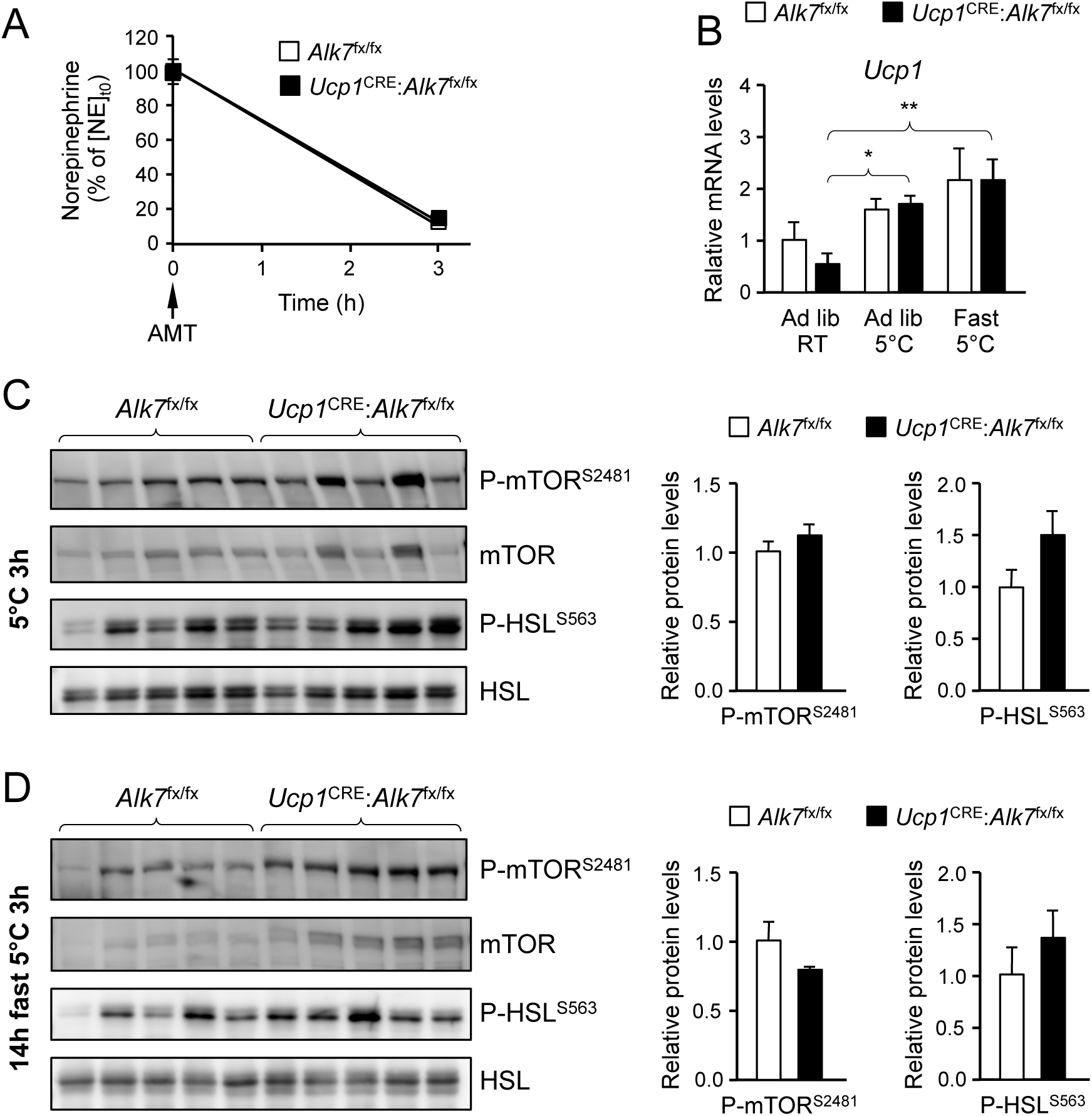
Unchanged norepinephirene signaling in iBAT of *Ucp1*^CRE^:*Alk7*^fx/fx^ mutant mice after fasting and acute cold exposure. (A) Norepinephrine (NE) turnover in iBAT. After 14h fasting (t=0), mice were injected with α-Methyl-DL-tyrosine methyl ester hydrochloride (AMT) and placed at 5° for 3h, after which NE was measured in iBAT. NE levels before AMT injection and cold exposure was normalized to 100%. Shown are averages ± SEM. N=6 mice per group. (B) Relative levels of *Ucp1* mRNA in iBAT of mutant and control mice assessed by Q-PCR. Values are presented relative to levels in control mice kept ad libitum at RT. Ad lib, mice fed *ad libitum* at room temperature (RT) or 3h at 5°C. The third group was fasted for 14h prior to 3h exposure at 5°C. Values are average ± SEM. N=5 mice per group. *, P<0.05; **, P<0.01. (C, D) Levels of mTORC2 phosphorylated at Ser^2481^ and HSL phosphorylated at Ser^563^ assessed by Western blotting of iBAT lystaes from control and mutant mice exposed to 5°C for 3h after *ad libitum* feeding (C) or after 14h fasting (D). Histogrmas show averages ± SEM of ooptical densities normalized to levels in control mice. N=5 mice per group.

**Supplementary Figure 10.**
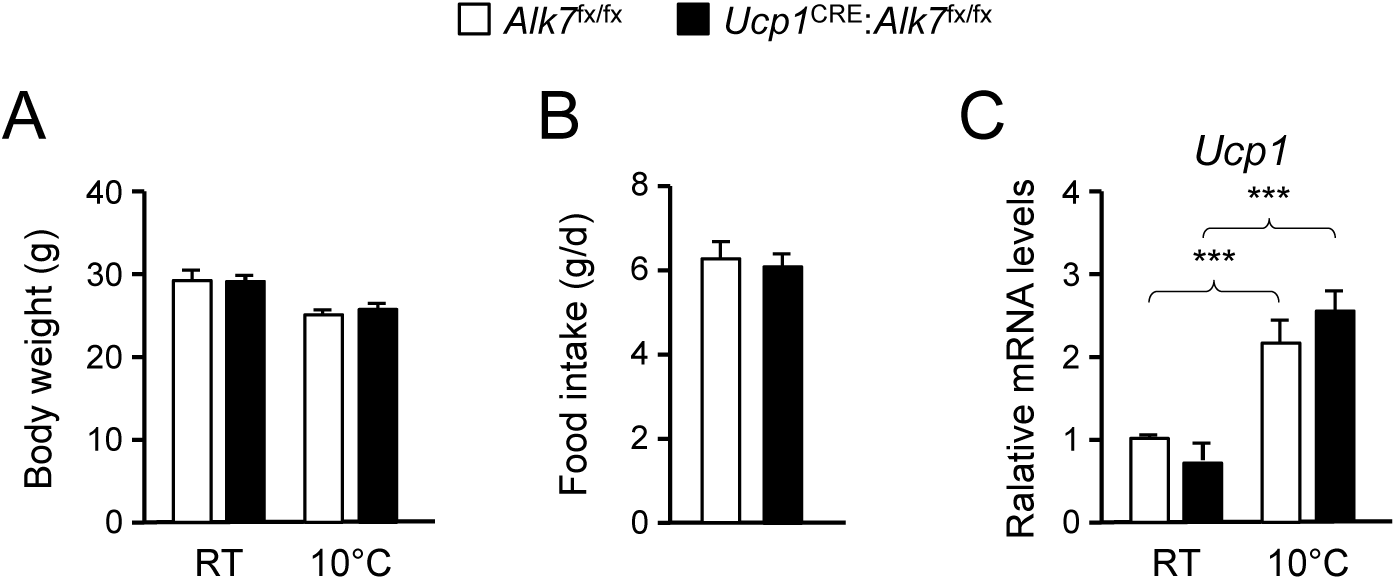
Body weight, food intake and *Ucp1* mRNA levels after chronic cold exposure (21 d at 10°C) in control and mutant mice. (A) Body weight. RT, room temperature. Shown are averages ± SEM. N=5 mice per group. (B) Daily food intake. Ahown are averages ± SEM. N=5 mice per group. (C) Levels of *Ucp1* mRNA in iBAT assessed by Q-PCR. Shown are averages relative to control levels ± SEM. Values were normalized to control mice at RT. N=5 mice per group. ***, P<0.001; one-way ANOVA with Tukey’s post test.

**Supplementary Figure 11.**
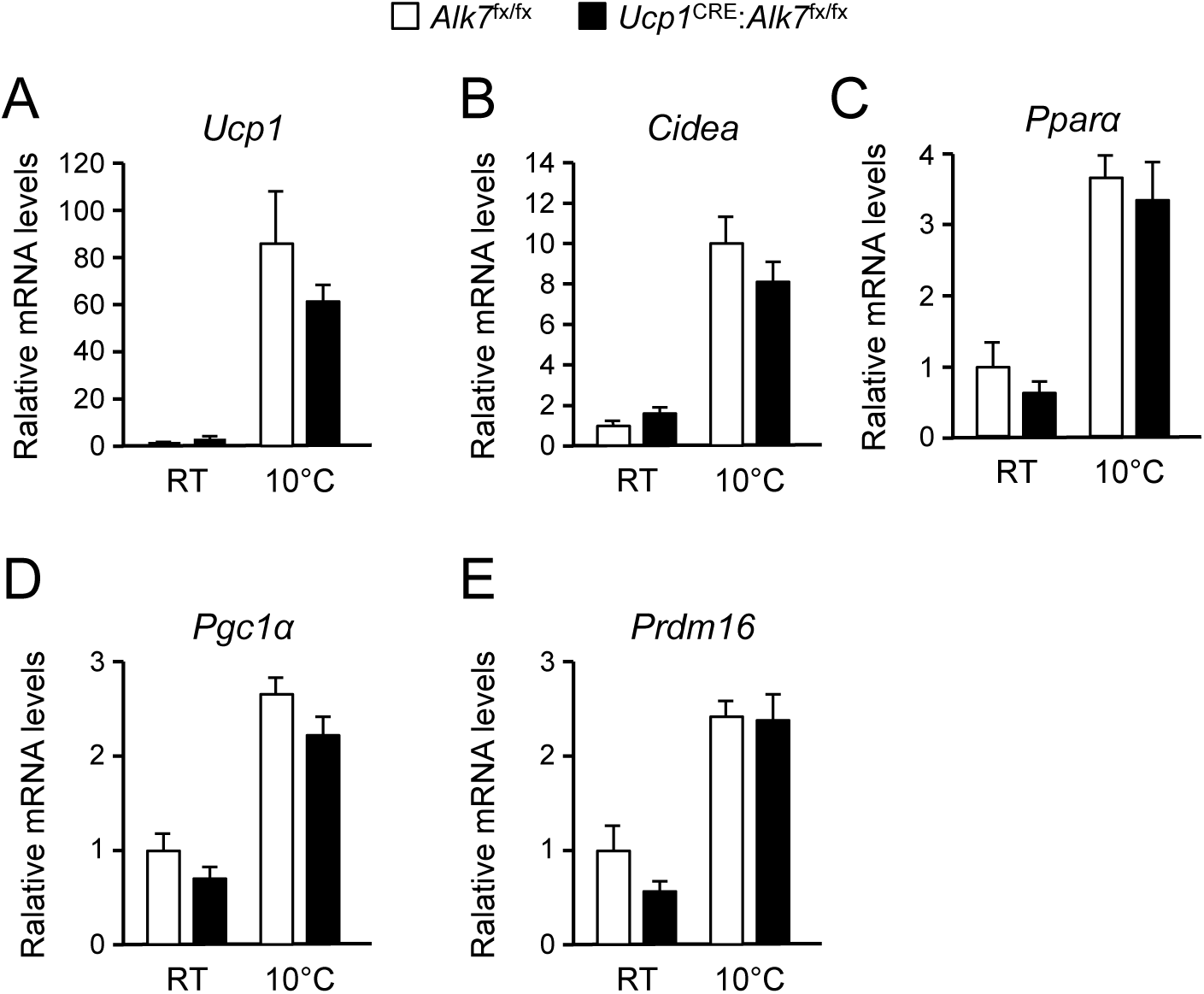
Expression of mRNA levels encoding thermogenic markers in inguinal WAT after chronic cold exposure (21 d at 10°C) in control and conditional mutant mice. (A to E) Levels of *Ucp1, Cidea, Pparα, Pgc1α* and *Prdm16* mRNAs in inguinal WAT assessed by Q-PCR in mice kept at room teperature and after chronic cold exposure. Shown are averages relative to control levels ± SEM. Values were normalized to contorl mice at RT. N=5 mice per group. ***, P<0.001; one-way ANOVA with Tukey’s post test.

**Supplementary Table 1:**
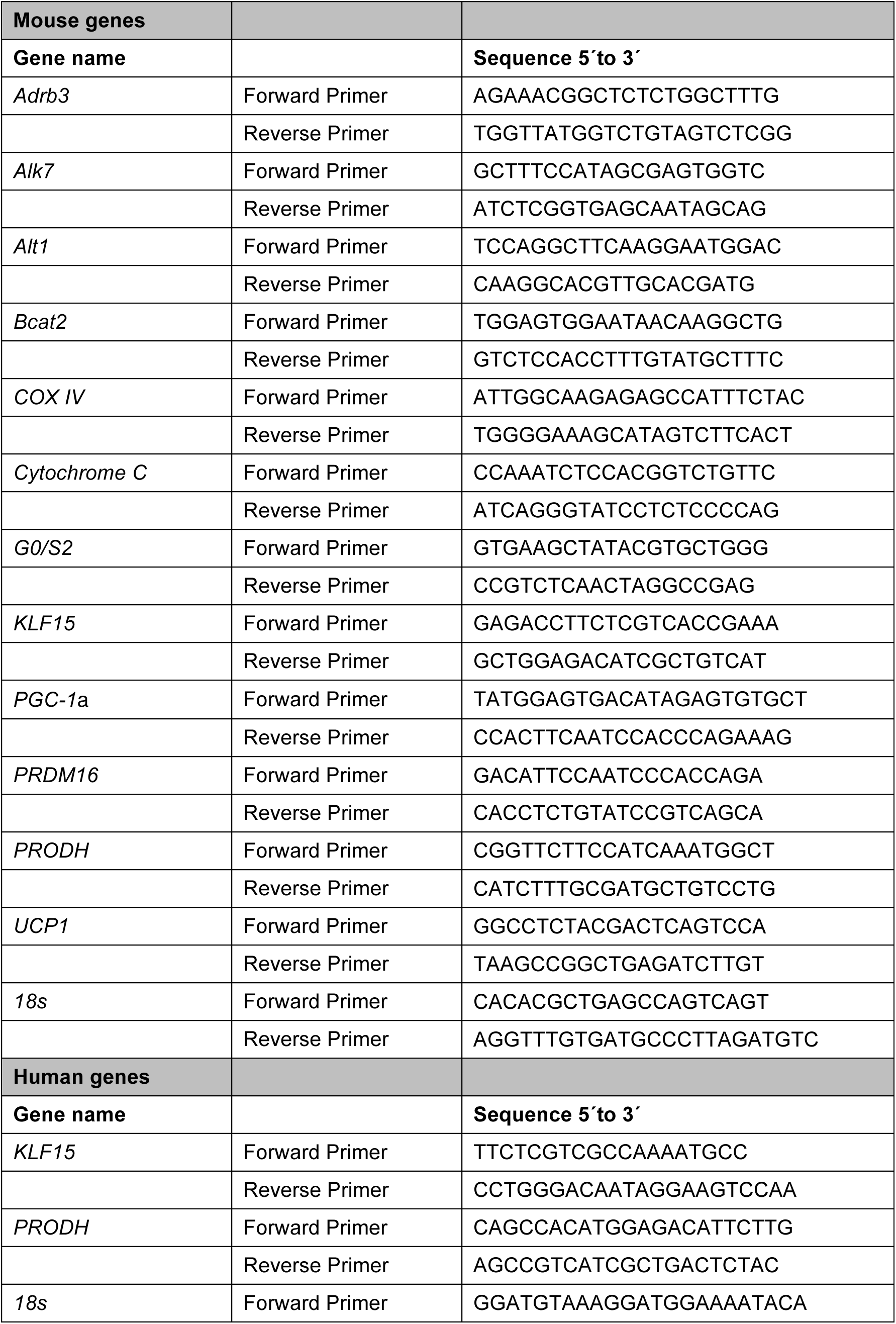
PCR primers.

**Supplementary table 2:** Antibodies.

| **Antibody** | **Reference** |
| --- | --- |
| Actin | Cell Signaling 4967 |
| Akt | Cell Signaling 9272 |
| ATGL | Cell Signaling 2138 |
| ATPB | Abcam 14730 |
| COX IV | Cell Signaling 4844 |
| HSL | Cell Signaling 4107 |
| mTOR | Cell Signaling 2972 |
| NDUFA10 | Santa Cruz 376357 |
| p-Akt<sub>473</sub> | Cell Signaling 9271 |
| p-HSL<sub>563</sub> | Cell Signaling 4139 |
| p-mTOR<sub>2448</sub> | Cell Signaling 2974 |
| p-mTOR<sub>2481</sub> | Cell Signaling 2971 |
| POX | Abcam 93210 |
| Rieske FeS | Santa Cruz 271609 |
| SDHA | Cell Signaling 5839 |
| UCP1 | Chemicon 1426 |

## References

Albert V, Svensson K, Shimobayashi M, Colombi M, Munoz S, Jimenez V, Handschin C, Bosch F, Hall MN (2016) mTORC2 sustains thermogenesis via Akt-induced glucose uptake and glycolysis in brown adipose tissue. EMBO Mol Med 8:232–246.

Andersson O, Korach-Andre M, Reissmann E, Ibáñez CF, Bertolino P (2008) Growth/differentiation factor 3 signals through ALK7 and regulates accumulation of adipose tissue and diet-induced obesity. Proc Natl Acad Sci USA 105:7252–7256.

Bertolino P, Holmberg R, Reissmann E, Andersson O, Berggren P-O, Ibáñez CF (2008) Activin B receptor ALK7 is a negative regulator of pancreatic beta-cell function. Proc Natl Acad Sci USA 105:7246–7251.

Brasaemle DL, Wolins NE (2016) Isolation of Lipid Droplets from Cells by Density Gradient Centrifugation. Current Protocols in Cell Biology 72:3.15.1–3.15.13.

Cannon B, Nedergaard J (2004) Brown adipose tissue: function and physiological significance. Physiol Rev 84:277–359.

Donald SP, Sun XY, Hu CA, Yu J, Mei JM, Valle D, Phang JM (2001) Proline oxidase, encoded by p53-induced gene-6, catalyzes the generation of proline-dependent reactive oxygen species. Cancer Res 61:1810–1815.

Emdin CA et al. (2019) DNA Sequence Variation in ACVR1C Encoding the Activin Receptor-Like Kinase 7 Influences Body Fat Distribution and Protects Against Type 2 Diabetes. Diabetes 68:226–234.

Fedorenko A, Lishko PV, Kirichok Y (2012) Mechanism of fatty-acid-dependent UCP1 uncoupling in brown fat mitochondria. Cell 151:400–413.

Frontini A, Cinti S (2010) Distribution and development of brown adipocytes in the murine and human adipose organ. Cell Metab 11:253–256.

Geiser F, Currie SE, O’Shea KA, Hiebert SM (2014) Torpor and hypothermia: reversed hysteresis of metabolic rate and body temperature. Am J Physiol Regul Integr Comp Physiol 307:R1324–R1329.

Goncalves RLS, Rothschild DE, Quinlan CL, Scott GK, Benz CC, Brand MD (2014) Sources of superoxide/H2O2 during mitochondrial proline oxidation. Redox Biol 2:901–909.

Gray S, Wang B, Orihuela Y, Hong E-G, Fisch S, Haldar S, Cline GW, Kim JK, Peroni OD, Kahn BB, Jain MK (2007) Regulation of gluconeogenesis by Krüppel-like factor 15. Cell Metab 5:305–312.

Guo T, Marmol P, Moliner A, Björnholm M, Zhang C, Shokat KM, Ibáñez CF (2014) Adipocyte ALK7 links nutrient overload to catecholamine resistance in obesity. Elife 3:e03245.

Haldar SM et al. (2012) Kruppel-like factor 15 regulates skeletal muscle lipid flux and exercise adaptation. Proc Natl Acad Sci U S A 109:6739–6744.

Hancock CN, Liu W, Alvord WG, Phang JM (2016) Co-regulation of mitochondrial respiration by proline dehydrogenase/oxidase and succinate. Amino Acids 48:859– 872.

Hao Q, Yadav R, Basse AL, Petersen S, Sonne SB, Rasmussen S, Zhu Q, Lu Z, Wang J, Audouze K, Gupta R, Madsen L, Kristiansen K, Hansen JB (2015) Transcriptome profiling of brown adipose tissue during cold exposure reveals extensive regulation of glucose metabolism. Am J Physiol Endocrinol Metab 308:E380–E392.

Inman GJ, Nicolás FJ, Hill CS (2002) Nucleocytoplasmic shuttling of Smads 2, 3, and 4 permits sensing of TGF-beta receptor activity. Mol Cell 10:283–294.

Jörnvall H, Reissmann E, Andersson O, Mehrkash M, Ibáñez CF (2004) ALK7, a receptor for nodal, is dispensable for embryogenesis and left-right patterning in the mouse. Mol Cell Biol 24:9383–9389.

Justice AE et al. (2019) Protein-coding variants implicate novel genes related to lipid homeostasis contributing to body-fat distribution. Nat Genet 51:452–469.

Kong X, Banks A, Liu T, Kazak L, Rao RR, Cohen P, Wang X, Yu S, Lo JC, Tseng Y-H, Cypess AM, Xue R, Kleiner S, Kang S, Spiegelman BM, Rosen ED (2014) IRF4 is a key thermogenic transcriptional partner of PGC-1α. Cell 158:69–83.

Korge P, Honda HM, Weiss JN (2003) Effects of fatty acids in isolated mitochondria: implications for ischemic injury and cardioprotection. Am J Physiol Heart Circ Physiol 285:H259–H269.

Matoba K, Lu Y, Zhang R, Chen ER, Sangwung P, Wang B, Prosdocimo DA, Jain MK (2017) Adipose KLF15 Controls Lipid Handling to Adapt to Nutrient Availability. Cell Rep 21:3129–3140.

Nedergaard J, Cannon B (2014) The browning of white adipose tissue: some burning issues. Cell Metab 20:396–407.

Nicholls DG, Locke RM (1984) Thermogenic mechanisms in brown fat. Physiol Rev 64:1–64.

Oelkrug R, Heldmaier G, Meyer CW (2011) Torpor patterns, arousal rates, and temporal organization of torpor entry in wildtype and UCP1-ablated mice. J Comp Physiol B, Biochem Syst Environ Physiol 181:137–145.

Olivares O et al. (2017) Collagen-derived proline promotes pancreatic ductal adenocarcinoma cell survival under nutrient limited conditions. Nat Commun 8:16031.

Pandhare J, Donald SP, Cooper SK, Phang JM (2009) Regulation and function of proline oxidase under nutrient stress. J Cell Biochem 107:759–768.

Phang JM (2019) Proline Metabolism in Cell Regulation and Cancer Biology: Recent Advances and Hypotheses. Antioxid Redox Signal 30:635–649.

Ramage LE, Akyol M, Fletcher AM, Forsythe J, Nixon M, Carter RN, van Beek EJR, Morton NM, Walker BR, Stimson RH (2016) Glucocorticoids Acutely Increase Brown Adipose Tissue Activity in Humans, Revealing Species-Specific Differences in UCP-1 Regulation. Cell Metab 24:130–141.

Reissmann E, Jörnvall H, Blokzijl A, Blokzijl A, Andersson O, Chang C, Minchiotti G, Persico MG, Ibáñez CF, Brivanlou AH (2001) The orphan receptor ALK7 and the Activin receptor ALK4 mediate signaling by Nodal proteins during vertebrate development. Genes Dev 15:2010–2022.

Rosen ED, Spiegelman BM (2014) What We Talk About When We Talk About Fat. Cell 156:20–44.

Rydén M, Imamura T, Jörnvall H, Belluardo N, Neveu I, Trupp M, Okadome T, Dijke ten P, Ibáñez CF (1996) A novel type I receptor serine-threonine kinase predominantly expressed in the adult central nervous system. J Biol Chem 271:30603–30609.

Sandoval-Guzmán T, Göngrich C, Moliner A, Guo T, Wu H, Broberger C, Ibáñez CF (2012) Neuroendocrine control of female reproductive function by the activin receptor ALK7. FASEB J 26:4966–4976.

Sasse SK, Mailloux CM, Barczak AJ, Wang Q, Altonsy MO, Jain MK, Haldar SM, Gerber AN (2013) The glucocorticoid receptor and KLF15 regulate gene expression dynamics and integrate signals through feed-forward circuitry. Mol Cell Biol 33:2104– 2115.

Schreiber R, Diwoky C, Schoiswohl G, Feiler U, Wongsiriroj N, Abdellatif M, Kolb D, Hoeks J, Kershaw EE, Sedej S, Schrauwen P, Haemmerle G, Zechner R (2017) Cold-Induced Thermogenesis Depends on ATGL-Mediated Lipolysis in Cardiac Muscle, but Not Brown Adipose Tissue. Cell Metab 26:753–763.e757.

Shimizu N, Yoshikawa N, Ito N, Maruyama T, Suzuki Y, Takeda S-I, Nakae J, Tagata Y, Nishitani S, Takehana K, Sano M, Fukuda K, Suematsu M, Morimoto C, Tanaka H (2011) Crosstalk between glucocorticoid receptor and nutritional sensor mTOR in skeletal muscle. Cell Metab 13:170–182.

Shin H, Ma Y, Chanturiya T, Cao Q, Wang Y, Kadegowda AKG, Jackson R, Rumore D, Xue B, Shi H, Gavrilova O, Yu L (2017) Lipolysis in Brown Adipocytes Is Not Essential for Cold-Induced Thermogenesis in Mice. Cell Metab 26:764–777.e765.

Shinoda K, Luijten IHN, Hasegawa Y, Hong H, Sonne SB, Kim M, Xue R, Chondronikola M, Cypess AM, Tseng Y-H, Nedergaard J, Sidossis LS, Kajimura S (2015) Genetic and functional characterization of clonally derived adult human brown adipocytes. Nat Med 21:389–394.

Villarroya F, Vidal-Puig A (2013) Beyond the sympathetic tone: the new brown fat activators. Cell Metab 17:638–643.

Wang W, Seale P (2016) Control of brown and beige fat development. Nat Rev Mol Cell Biol 17:691–702.

Wu J, Cohen P, Spiegelman BM (2013) Adaptive thermogenesis in adipocytes: is beige the new brown? Genes Dev 27:234–250.

Yamamoto J, Ikeda Y, Iguchi H, Fujino T, Tanaka T, Asaba H, Iwasaki S, Ioka RX, Kaneko IW, Magoori K, Takahashi S, Mori T, Sakaue H, Kodama T, Yanagisawa M, Yamamoto TT, Ito S, Sakai J (2004) A Kruppel-like factor KLF15 contributes fasting-induced transcriptional activation of mitochondrial acetyl-CoA synthetase gene AceCS2. J Biol Chem 279:16954–16962.

Yang X, Lu X, Lombès M, Rha GB, Chi Y-I, Guerin TM, Smart EJ, Liu J (2010) The G(0)/G(1) switch gene 2 regulates adipose lipolysis through association with adipose triglyceride lipase. Cell Metab 11:194–205.

